# Nek8445, a protein kinase required for microtubule regulation and cytokinesis in *Giardia lamblia*

**DOI:** 10.1101/719005

**Authors:** Kelly M. Hennessey, Germain C.M. Alas, Ilse Rogiers, Renyu Li, Ethan A. Merritt, Alexander R. Paredez

## Abstract

*Giardia* has 198 Nek kinases whereas humans have only 11. *Giardia* has a complex microtubule cytoskeleton that includes eight flagella and several unique microtubule arrays that are utilized for parasite attachment and facilitation of rapid mitosis and cytokinesis. The need to regulate these structures may explain the parallel expansion of the number of Nek family kinases. Here we use live and fixed cell imaging to uncover the role of Nek8445 in regulating *Giardia* cell division. We demonstrate that Nek8445 localization is cell cycle regulated and this kinase has a role in regulating overall microtubule organization. Nek8445 depletion results in short flagella, aberrant ventral disc organization, loss of the funis, defective axoneme exit and altered cell shape. The axoneme exit defect is specific to the caudal axonemes, which exit from the posterior of the cell, and this defect correlates with rounding of the cell posterior and loss of the funis. Our findings implicate a role for the funis in establishing *Giardia’s* cell shape and guiding axoneme docking. On a broader scale our results support the emerging view that Nek family kinases have a general role in regulating microtubule organization.

## Introduction

The protozoan parasite *Giardia lamblia (synonymous with G. intestinalis and G. duodenalis)* infects more than 200 million people each year (Lane and Lloyd, 2002; Lalle, 2010; Ansell *et al.*, 2015). The divergence of parasite protein kinases from their homologs in humans has been identified as an opportunity for the development of new therapeutic interventions (Rotella, 2012). The *Giardia* kinome (strain WB) contains 278 protein kinases, 80 of which constitute the core kinome from 49 different classes of kinases while the remainder are all Nek Kinase homologs (Manning *et al.*, 2011). The highly expanded *Giardia* Nek family contains 198 kinases, compared to only one in budding yeast, seven in *Arabidopsis,* 11 in humans, 13 in *Chlamydomonas, and 39 in Tetrahymena* (Wloga *et al.*, 2006; Manning *et al.*, 2011; Takatani *et al.*, 2015).

Nek kinases are named for their homology to the *Aspergillus nidulans* NIMA (never in mitosis A) kinase. NIMA related kinases (Nek or NRK) kinases have been shown to control mitotic entry, centrosome separation, and flagella length in other eukaryotes (Faragher and Fry, 2003; O’Connell *et al.*, 2003; Bradley and Quarmby, 2005; Wloga *et al.*, 2006; Chen and Gubbels, 2013), but may also have a more general role in regulating microtubule organization (Takatani *et al.*, 2015). The number of Nek kinases is typically expanded in the genomes of eukaryotes that build cilia or flagella (Parker *et al.*, 2007). Notably, *Giardia* possesses four pairs of flagella that undergo a complex developmental cycle (Nohynkova *et al.*, 2006) (See Figure 1 for diagram of *Giardia* structures). As pairs of flagella do not share paired basal bodies, maintaining the identity of each flagella is complex and the regulatory mechanism remains uncharacterized (Nohynkova *et al.*, 2006). The complexity of maintaining eight flagella identities could explain the development of a greatly expanded set of Nek kinases in *Giardia.*

**Figure 1.**
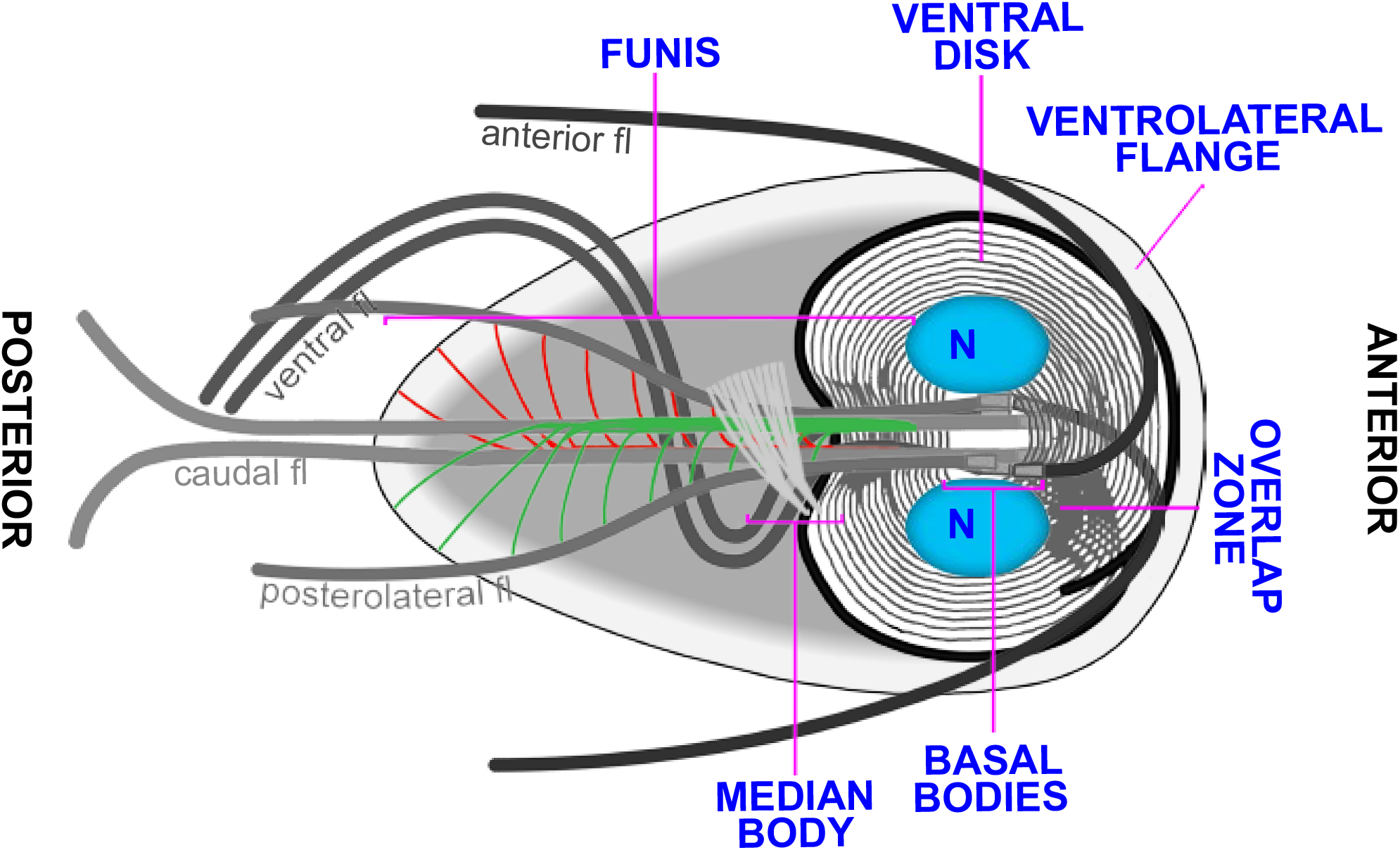
Diagram of *Giardia* emphasizing microtubule-based structures. The microtubule cytoskeleton includes the ventral disc, median body, funis, anterior, posterolateral, ventral and caudal flagella sets.

Flagella positioning and identity are essential for *Giardia* cellular processes including pathogenicity. Unlike most other eukaryotes that dock their basal bodies just under the plasma membrane, *Giardia* positions its eight basal bodies between the two nuclei in the center of the cell. As a result, a significant portion of each axoneme runs through the cytoplasm of the cell before it exits and is sheathed by membrane (McInally and Dawson, 2016; McInally *et al.*, 2019b). We use the term flagella to refer to the membrane bound portion of axonemes while the portion of the axonemes running through the cell body are referred to as cytoplasmic axonemes. *Giardia* cytoplasmic axonemes serve as tracks for cargo trafficking and have also been implicated in force generation for cell division (Hardin *et al.*, 2017). Each of *Giardia’s* flagella pairs beat with unique waveforms, further indicating the complexity of specifying flagella identity and function (Lenaghan *et al.*, 2011). For example, the caudal axoneme pair, which runs the length of the cell body, is thought to flex the entire body of the cell to contribute to swimming via a flipper type mechanism (Lenaghan *et al.*, 2011). The anterior and posterolateral flagella generate power strokes for motility; while the ventral flagella, with a unique fin-like appendage, beat in a sinusoidal waveform thought to generate fluid flows beneath and around *Giardia* trophozoites (Holberton, 1973; Lenaghan *et al.*, 2011; Halpern *et al.*, 2017).

In addition to four unique sets of flagella, *Giardia* has several novel microtubule structures including the ventral adhesive disc, median body, and funis (also known as the caudal complex/ With the exception of the median body, each of these structures is associated with specific sets of basal bodies (McInally and Dawson, 2016). The ventral disc is a complex dome shaped structure composed of approximately 100 microtubules in a one and a quarter turn helical sheet and is thought to mediate attachment through a suction cup like mechanism (Erlandsen *et al.*, 2004; Hansen *et al.*, 2006; Brown *et al.*, 2016; Nosala *et al.*, 2018). The funis is a pair of microtubule sheets that originate at the basal bodies. One sheet runs above the cytoplasmic caudal axoneme pair while the other runs below the caudal cytoplasmic axonemes. Toward the posterior of the cell, the funis microtubules separate from the main sheet to form rib-like structures that lock the cytoplasmic axonemes of the caudal and posterolateral flagella together (Benchimol *et al.*, 2004; Carvalho and Monteiro-Leal, 2004). The median body is a reservoir of microtubules and associated binding proteins used to support *Giardia’s* rapid mitosis (Hardin *et al.*, 2017; Horlock-Roberts *et al.*, 2017). Remarkably, mitosis is completed in approximately 7 minutes and this includes duplicating axonemes and building two new ventral discs (Hardin *et al.*, 2017). Currently, little is known about how the assembly and disassembly dynamics of these structures are regulated. As *Giardia* lacks an anaphase promoting complex, it has been suggested that kinases may play an enhanced role in regulating mitotic events (Gourguechon *et al.*, 2013). Indeed, Nek kinases have been implicated in the regulated assembly and disassembly of microtubule based structures during the cell cycle (Smith *et al.*, 2012).

The *Giardia* Nek kinase homologs with the highest sequence identity for human Nek1 and Nek2 were previously studied (Smith *et al.*, 2012). The Nek1 homolog *Gl*Nek1 localized to the ventral disc while the Nek2 homolog *G/*Nek2 localized to the cytoplasmic axonemes of the anterior flagella. Both kinases re-organize during mitosis and over-expression studies showed that *Gl*Nek1 has a role in ventral disc disassembly while *Gl*Nek2 over expression interfered with excystation (hatching from the cyst form). Other *Giardia* Nek kinases have not yet been individually studied.

Here, we characterize the role of Nek kinase GL50803_8445 (Nek8445) in *Giardia* cell proliferation. Nek8445 has direct orthologs (>90% sequence identity) in other *Giardia* assemblages (strains), but exhibits only 30-40% sequence identity to other Nek paralogs in the expanded *Giardia* Nek kinome. This is consistent with a functional role in *Giardia* that is both essential and distinct from that of other Nek family kinases. We previously identified this kinase as a potential therapeutic target due to the presence of a small gatekeeper residue, which makes it susceptible to a specific class of kinase inhibitors with relatively low activity against human kinases (Hennessey *et al.*, 2016). These “bumped kinase inhibitors” (BKIs) exploit the property that most mammalian protein kinases have a large gatekeeper residue that limits accessibility to a hydrophobic pocket in the active site. Various BKIs have been shown to selectively and effectively block kinase-associated processes in the life cycle of parasites such as *Toxoplasma gondii, Cryptosporidia parvum* and *Plasmodium falciparum* (Keyloun *et al.*, 2014). In *Giardia* there are at least eight small gatekeeper kinases (Hennessey *et al.*, 2016) Although we identified inhibitors that target Nek8445 they also target additional small gatekeeper kinases, which currently limits their usefulness in uncovering the role of Nek8445.

We previously determined that Nek8445 depleted cells are severely impeded in growth, often abnormally multinucleated, and unable to maintain attachment (Hennessey *et al.*, 2016). The initial study focused on defects at 48 hours after morpholino knockdown, which corresponded with the endpoint of our growth assays. By this time cellular organization was severely disrupted and therefore the primary defect was not apparent. However, the cytokinesis and attachment defects are potentially explained by aberrant flagella function. Unlike other eukaryotes that use an actomyosin contractile ring to guide membrane delivery and drive cytokinesis, *Giardia* utilizes nascent cytoplasmic axonemes to coordinate membrane trafficking into the furrow while the mature flagella propel daughter cells apart for abscission (Hardin *et al.*, 2017). The flagella may also have a role in parasite attachment (Holberton, 1973). It has been proposed that flows generated by the ventral flagella transit through gates in the ventral disc which can close off to lock down attachment (Dawson and Nosala, 2017). Here we examine the role of Nek8445 in *Giardia’s* unique mode of cytokinesis.

## Methods & Materials

### Parasite culture

*Giardia lamblia* wild-type strain *WBC6* (ATCC 50803) cells were grown in TYI-S-33 medium supplemented with 10% bovine serum and 0.05 mg/mL bovine bile (Keister, 1983). Cells were cultured at 37° under hypoxic conditions using 15 mL polystyrene screw-cap tubes (Corning Life Sciences DL).

### Vector construction and transfection

With the exception of Rab11, the constructs used in this study were C-terminal fusions. All constructs were integrated into the genome using single site recombination to maintain endogenous expression levels (Gourguechon and Cande, 2011). Primer sequences and restriction enzymes are shown in Table S1. All PCR amplifications were performed using iProof DNA polymerase (Bio-Rad). Typically, an amplicon of ~ 1 kb in length that lacked the start codon was cloned in frame to a C-terminal triple-hemagglutinin epitope tag (3xHA) of the pKS_3HA_Neo or PAC plasmid (Gourguechon and Cande, 2011). The plasmids were linearized at a unique restriction site in the coding region of the gene of interest and ~5 μg of DNA was used to transfect wild-type *Giardia.* Transfected cells were selected with G418 at 40 μg/mL or puromycin at 45 μg/mL.

To follow Nek8445 localization throughout mitosis and cytokinesis, the C terminus of GL50803_8445 was tagged with an 11-amino acid flexible linker and the fluorescent protein mNeonGreen (Allele Biotechnology). The mNG-N11-Neo vector was constructed for this purpose by amplifying mNeonGreen using the primers listed in Table S1. The PCR product was cloned into the pKS 3HA Neo vector (Gourguechon and Cande, 2011) after excising the 3HA tag with BamH1 and EcoR1.

### Live cell imaging

Cells were imaged in 35mm glass bottom dishes overlaid with agarose growth medium at 37° C under hypoxic conditions as described in (Hardin *et al.*, 2017).

### Immunofluorescence microscopy analysis

Fixed cell imaging was performed as described in (Paredez *et al.*, 2011). In cases where a higher mitotic index was desired, the starve and release method was utilized before fixation (Sagolla *et al.*, 2006). Briefly, four-day-old past confluent cultures were inverted to loosen dead cells and then the dead cells and spent medium were poured off and replaced with fresh medium. Four hours later, the cells were pelleted at 500xg at room temperature. The pellet and remaining attached cells were fixed in PME (100 mM Pipes pH 7.0, 5 mM EGTA, 10 mM MgSO4) plus 2% paraformaldehyde, 0.025% Triton X-100, 100 μM MBS, and 100 μM EGS for 30 minutes at 37°C. Cells were again pelleted, washed with PME, resuspended with PME, and adhered to poly-L-lysine (Sigma-Aldrich) coated cover-slips. Cells were permeabilized in PME + 0.1% Triton X-100 for 10 minutes then washed 2X with PME + 0.05% Triton X-100 and blocked for 30 minutes in PMEBALG (PME + 1% BSA, 0.1% NaN3, 100 mM lysine, 0.5% cold water fish skin gelatin (Sigma Aldrich, St. Louis, MO).

Cells were stained with rabbit anti-*Gl*Actin antibody 28PB+1 (Paredez *et al.*, 2011) and mouse monoclonal anti-HA (Clone HA7, Sigma-Aldrich) both diluted 1:125 in PMEBALG and incubated over-night. After three subsequent washes with PME + 0.05% Triton X-100, cells were incubated with Alexa-488 goat anti-mouse and Alexa-555 goat anti-α-rabbit (Sigma-Aldrich, St. Louis, MO) (diluted 1:150 in PMEBALG) for 1 h. Cells were washed three times with PME + 0.05% Triton X-100. The coverslips were mounted with ProLong Gold or Prolong Diamond anti-fade plus DAPI, (ThermoFisher Scientific, Rockford, IL). Fluorescence deconvolution microscopy images were collected as described (19). For all standard resolution microscopy experiments a minimum of 200 cells were examined with at least 20 imaged per experiment. Structured illumination images were acquired on an OMX SR (GE Healthcare).

### Morpholino knockdown and growth assays

Trophozoites were cultured until 85-100% confluence, placed on ice for 30 minutes to detach cells, pelleted at 500xg for 5 minutes, and medium was replaced with 1.0 mL fresh ice cold *Giardia* growth medium. Cells and cuvettes were chilled on ice. Lyophilized morpholinos listed in Table S1 (Gene Tools, LLC, Philomath, OR) were resuspended in sterile water and 30 μL of a 100 mM morpholino stock was added to 300 μL of cells in a 4 mm cuvette. We used Gene Tools, LLC standard control morpholino as a negative control. Cells were electroporated (375V, 1000 μF, 750 Ohms, GenePulser Xcell, Bio-Rad, Hercules, CA) and returned to growth medium. See (Krtkova and Paredez, 2017) for a detailed protocol.

### Western blot analysis

*Giardia* trophozoites were chilled on ice for 30 minutes to detach them from the culture tube side walls. Cells were pelleted at 700xg, washed once in HBS (HEPES buffered saline), then resuspended in 300 μL of lysis buffer (50 mM Tris pH 7.5, 150 mM NaCl, 7.5% Glycerol, 0.25 mM CaCl2, 0.5 mM DTT, 0.5 mM PMSF, 0.1% Triton X-100, Halt 100X Protease Inhibitor Cocktail (ThermoFisher Scientific), then sonicated. The lysate was cleared by centrifugation at 10,000xg for 10 minutes at 4°C and then boiled in 2x Laemmli Sample Buffer (Bio-Rad). After SDS-PAGE, samples were transferred to PVDF membrane (Immobilon-FL) following the manufacturers’ directions. Primary polyclonal rabbit anti-*Gl*Actin 28PB+1 (21), monoclonal anti-HA mouse HA7 antibodies (IgG1; Sigma-Aldrich), 6-11B-1 anti-acetylated tubulin (IgG2b), DMA1 anti-alpha Tubulin (IgG1, Sigma-Aldrich), were diluted 1:2500 in blocking solution (5% dry milk, 0.05% Tween-20 in TBS). Secondary isotype specific anti-mouse Alexa-555 or anti-rabbit Alexa-647 antibodies (Molecular Probes) were used at 1:2500. Horseradish peroxidase-linked anti-mouse or anti-rabbit antibodies (Bio-Rad) were used at 1:7,000. Multiplexed immunoblots were imaged on a Chemidoc MP (Bio-Rad). Digital images were exported to 16 bit TIFF files and protein levels were determined by measuring integrated density using ImageJ (Schneider *et al.*, 2012).

Protein levels in Figure 2A were normalized using actin as a loading control. For equal loading in Figure 4D the cells were counted using a MoxiZ coulter counter (ORFLO) and the total cell number was adjusted to match the tube with the lowest cell number before making lysates. Reported values are the mean of three independent experiments.

**Figure 2.**
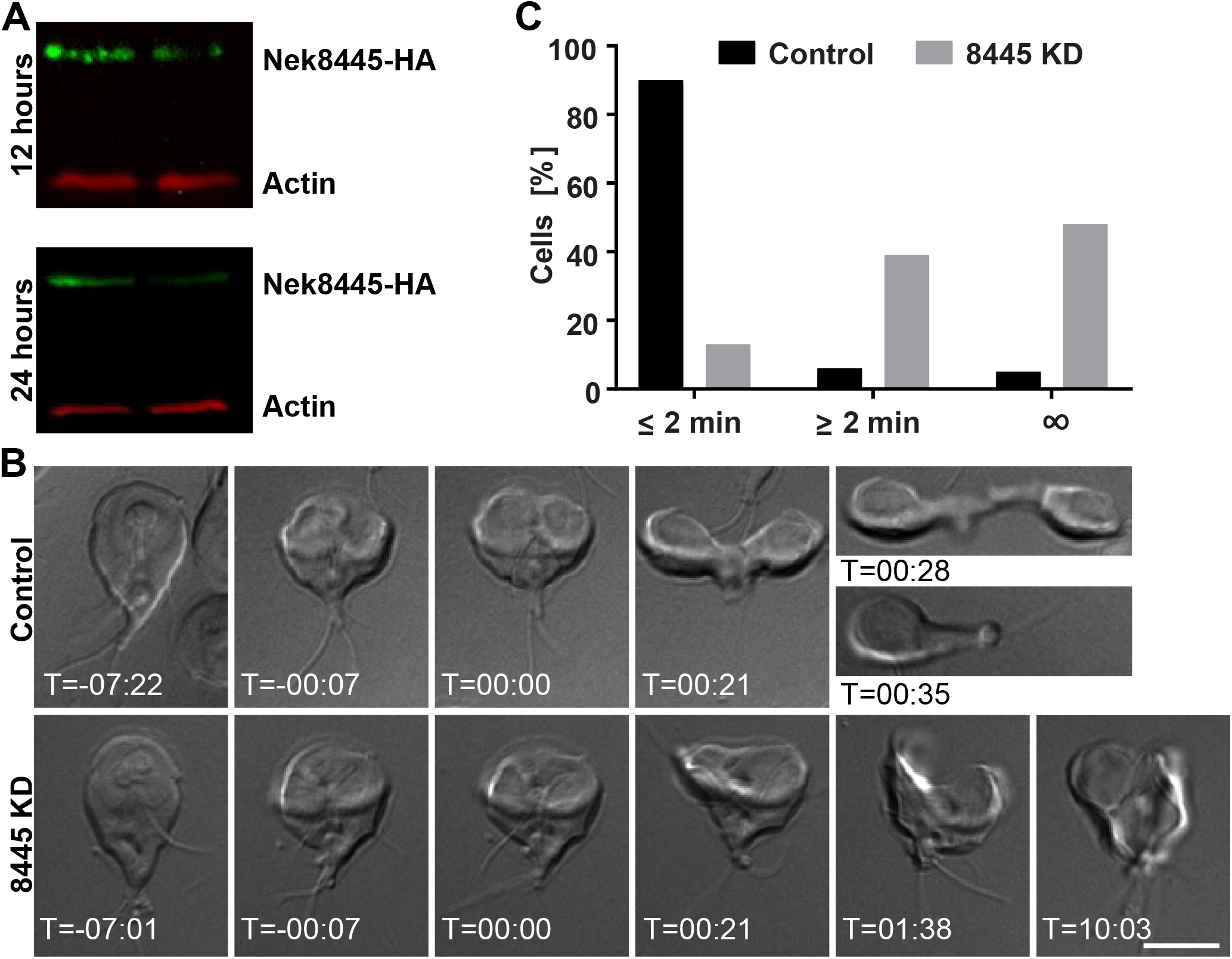
Nek8445 is required for cytokinesis. (A) Multiplex immunoblot showing Nek8445-HA depletion 12 and 24 hours after Nek8445 antisense and control morpholino treatment. (B) Representative still images from a 4D time-lapse movie showing failed cytokinesis in Nek8445 depleted cells (see Video S1). T=0 set to the initiation of cytokinesis, which occurs ~7 minutes after the initiation of mitosis as indicated by anterior flagella re-organization. (C) Histogram of cytokinesis timing. Only 13% of Nek8445 depleted cells complete cytokinesis within 2 minutes versus 90% for the control treatment (n=88 Control Morpholino, n=56 Nek8445 KD). Scale bar = 5 μm.

### Scanning Electron Microscopy

For scanning electron microscopy (SEM) analysis, trophozoites were pelleted at 700xg for 7 minutes, fixed with 2.5% glutaraldehyde in 0.1 M phosphate buffered saline (PBS), pH 7.4 and incubated for 1 h. After fixation, cells were pelleted again at 700xg for 7 minutes and washed twice and resuspended in PBS. Cells were adhered to poly-L-lysine (Sigma-Aldrich) coated coverslips for 15 minutes then washed three times for 5 minutes with PBS and post-fixed in 1% OsO_4_ for 1 h. Cells were washed three times for 5 minutes with dH_2_O, dehydrated in alcohol, dried to critical point with CO_2_, sputter coated with gold and analyzed on JEOL JSM-840A SEM.

### Accession numbers

**Genes/proteins**

**Nek8445: GL50803_8445; Axoneme Exit Site Protein: GL50803_8855, Rab11: GL50803_1695, Actin: GL50803_40817; δ-giardin: GL50803_86676**

## Results

### Nek8445 function is required for cytokinesis

The initial observation that Nek8445 plays a role in cytokinesis was based on analysis of Nek8445 depleted cells fixed and analyzed 48 h after treatment with translation-blocking antisense morpholinos (Hennessey *et al.*, 2016). These cells were abnormally multinucleate (more than 2 nuclei) with a roughly corresponding increase in flagella number (Figure S1). Cellshape and flagella organization were disrupted in the multi-nucleate cells. Therefore, it was not clear if the cytokinesis defect was a primary consequence of Nek8445 knockdown or if disrupted cell polarity led to failed cytokinesis. We recently developed methods for long-term 4D DIC time-lapse imaging to follow *Giardia* trophozoites through the cell cycle (Hardin *et al.*, 2017). We sought to film Nek8445 depleted cells undergoing cytokinesis to test for changes in the timing of events and gain insight into the underlying cause of failed cytokinesis. We planned to examine Nek8445 depleted cells from 20 – 26 h after morpholino treatment as in our previous study of *Giardia* cytokinesis. However, the cells were poorly adherent and the majority of the attached cells had already failed cytokinesis or were abnormally organized (Figure S2). Therefore, we decided to film cells from 12-18 h after knockdown. The majority of cells at this time point displayed typical organization as assessed by DIC imaging (Figure S2), allowing us to follow cytokinesis in long-term 4D imaging experiments. Supporting significant depletion of Nek8445 at this time point, Western blotting indicated that antisense morpholino treatment reduced Nek8445 levels by 41% compared with cells treated with GeneTools LLC’s standard control morpholino at 12 hours post electroporation (~1.5 cell cycles after treatment; Figure 2A).

Upon the initiation of cytokinesis, the cleavage furrow often failed to progress (Figure 2B, Video S1). Under our assay conditions, 90% of control morpholino treated cells completed cytokinesis within two minutes (n=88) similar to our previous results (Hardin *et al.*, 2017). In contrast, only 13% of Nek8445 depleted cells (n=56) completed cytokinesis within two minutes. Additionally, 48% of Nek8445 depleted cells never completed cytokinesis compared to just 4.5% non-completion observed for cells treated with the control morpholino (Figure 2C). Although Nek kinases are implicated in centrosome duplication and regulation of mitosis, we did not observe any mitotic defects as indicated by typical nuclear segregation in Nek8445 depleted cells (Video S1). Remarkably, the cell division defect resulting from Nek8445 knockdown is stronger than the phenotype of any cytoskeleton or membrane trafficking component previously studied (Hardin *et al.*, 2017), suggesting a central regulatory role for Nek8445 in coordinating cytokinesis.

### Nek8445 localization is cell cycle regulated

We previously reported that Nek8445-3HA localized to the plasma membrane and nuclear envelope in interphase cells (Hennessey *et al.*, 2016). Many cell cycle regulated proteins, including NEK kinases, change their localization during mitosis (Mahjoub *et al.*, 2004; Smith *et al.*, 2012; Chen and Gubbels, 2013). To follow Nek8445 localization throughout the cell cycle in live cells, the C-terminus of Nek8445 was tagged with the green fluorescent protein, mNeonGreen (Shaner *et al.*, 2013), generating Nek8445-mNeonGreen (Nek8445-mNG). This construct was integrated into the genome to ensure endogenous expression levels (Gourguechon and Cande, 2011). At the initiation of mitosis, Nek8445-mNG translocated from the cell cortex to become concentrated around the two nuclei and along the cytoplasmic axonemes (Figure 3A; Video 2). Late in mitosis Nek8445-mNG also concentrate at what appear to be the tips of the cytoplasmic axonemes as the axonemes were being re-arranged (Video 2). During cytokinesis Nek8445-mNG remained in association with the nuclear envelopes as well as portions of the anterior, caudal, and posterolateral cytoplasmic axonemes.

**Figure 3.**
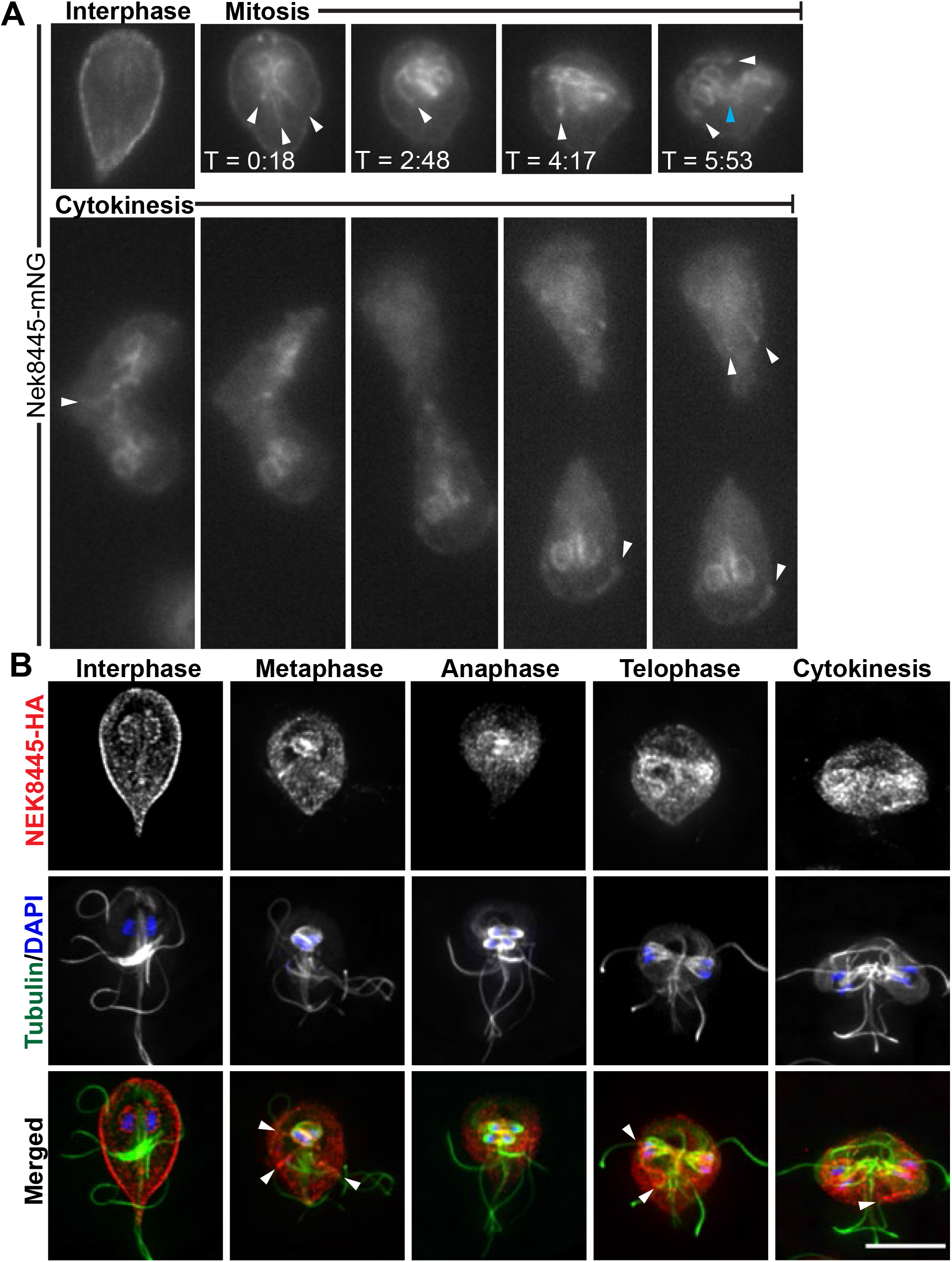
Nek8445 re-localizes during mitosis to associate with axoneme microtubules (A) Still images form time-lapse movie of Nek8445-mNG showing Nek8445 translocating from the cell cortex to the nuclear envelope, and axoneme microtubules (See Video S2). White arrowheads indicate Nek8445 in association with cytoplasmic axonemes, cyan arrowhead marks nascent axonemes pointing into the developing furrow. (B) Immunofluorescence analysis of Nek8445-HA (red), microtubules (6-11B-1 antibody, green), and nuclei (DAPI, blue) throughout the cell cycle. Scale bar = 5 μm.

To confirm association of Nek8445 with the cytoplasmic axonemes of flagella during mitosis and cytokinesis, we co-localized Nek8445-3HA and microtubules in fixed cells (Figure 3B). Indeed, Nek8445-3HA co-localized with the cytoplasmic axonemes of flagella including some association with the nascent axonemes that act as tracks for delivering membrane to the cleavage furrow (Hardin *et al.*, 2017). Overall the two chimeric proteins have similar localization, however Nek8445-3HA clearly associates with the nuclear envelope of interphase cells while Nek8445-mNG only weakly associated with the nuclei until the initiation of mitosis.

We previously established that Rab11 positive vesicles traffic along cytoplasmic axonemes and into the cleavage furrow to support furrow progression (Hardin *et al.*, 2017). The localization of Nek8445 to the cytoplasmic axonemes prompted us to ask if Nek8445 and Rab11 might have a functional relationship. Co-localization of HA-Rab11 with Nek8445-mNG revealed that while both proteins translocate from the plasma membrane to the intracytoplasmic axonemes during mitosis and cytokinesis, they were rarely seen to be on cytoplasmic axonemes at the same time (Figure S3). Therefore, it is unlikely that Nek8445 is directly regulating membrane trafficking associated with furrowing. Instead, Nek8445 may have a role in regulating axoneme/flagella function, which is required for force generation during cytokinesis (House *et al.*, 2011; Hardin *et al.*, 2017).

### Nek8445 depleted cells have short flagella and lack median bodies

The microtubule cytoskeleton plays a major role in establishing *Giardia’s* unique shape and in generating forces for driving cells apart during cytokinesis (Mariante *et al.*, 2005; Tumova *et al.*, 2007; Hardin *et al.*, 2017). Defects in microtubule regulation could account for defects in cell division (Mariante *et al.*, 2005). Therefore, we examined tubulin staining in Nek8445 depleted cells (Figure 4A). We observed that ~17% (n>300) of the Nek8445 depleted cells had noticeably short flagella and either round or oval-shaped cell bodies (Figure 4A and 4B).

**Figure 4.**
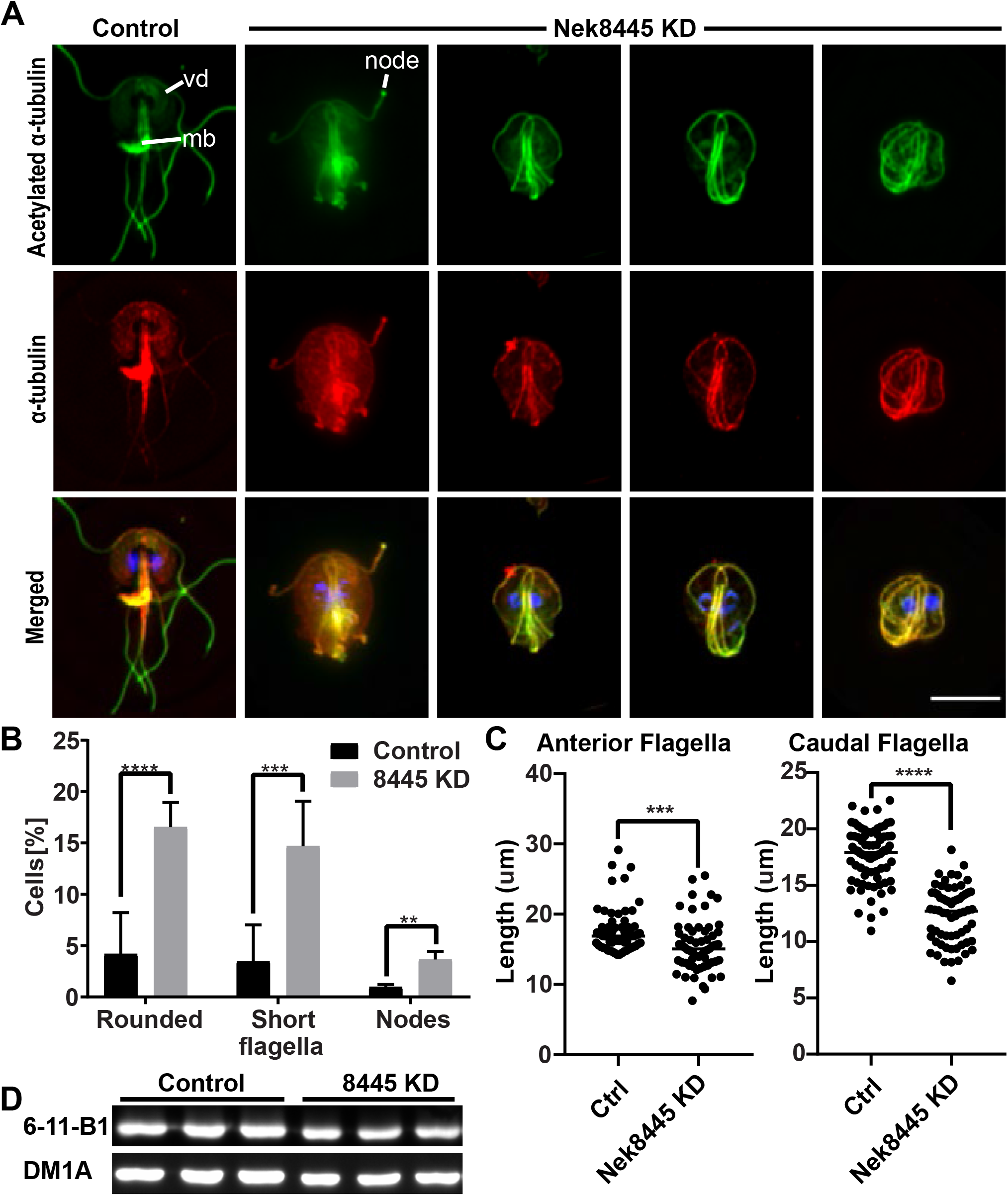
Nek8445 regulates microtubule organization. (A) Immunofluorescence staining of acetylated and total α-tubulin. Acetylated tubulin is stained with the anti-acetylated α-tubulin antibody 6-11B-1 (green) and total tubulin is stained with the anti-α-tubulin antibody DM1A (red). Note changes in overall tubulin staining after Nek8445 depletion including reduced ventral disc (vd) staining, the lack of a median body (mb) in rounded cells, and nodes of tubulin staining at the ends of flagella. Scale bar = 5 μm. (B) Quantification of mutant phenotypes observed after Nek8445 depletion (n>300 cells). (C) Measurement of anterior and caudal axoneme length from control morpholino and Nek8445 knockdown cells (n=76 measurements from control cells and n=64 measurements for 8445 knockdown cells). (D) Immunoblotting for total tubulin (DM1A) vs acetylated tubulin (6-11B-1 antibody) in Nek8445 and control morpholino treated cells. Lysates were prepared from equal numbers of cells for each replicate. There is no significant difference in the ratio of acetylated to total tubulin although overall tubulin levels are decreased by ~25% in Nek8445 depleted cells.

We measured the entire length of anterior and caudal axonemes from randomly selected control and Nek8445 translation blocking morpholino treated cells. The anterior axonemes were modestly shorter (Control Morpholino: 17.5 ± 0.34 μm, n=76; Nek8445 KD 15.5 ± 0.48 μm, n=60) while the caudal axonemes were 30% shorter (Control Morpholino: 17.7 ± 0.27 μm, n=80; Nek8445 KD: 12.3 ± 0.31 μm, n=64). The values reported here likely under-report the shortening defect as the knockdown cells with the most severe flagella shortening had tangled axonemes, which we omitted from our measurements because they were too convoluted to trace. Similarly, we did not attempt to measure the posterolateral or ventral axonemes as they were often difficult to identify, due to being mispositioned, in the Nek8445 knockdown cells.

We also noted that the rounded cells lacked median bodies (Figure 4A). Upon quantitation, we found that 46.8% (n=158) of all Nek8445 depleted cells lacked median bodies versus just under 15% (n=302) of the control morpholino treated cells. Note that this microtubule reservoir is consumed during mitosis and then grows in size as cells prepare for the next round of division (Hardin *et al.*, 2017; Horlock-Roberts *et al.*, 2017). Additionally, we identified abnormal accumulation of tubulin in nodes at the tips of flagella in the Nek8445 knockdown cell (Figure 4A and 4B). Short flagella and accumulation of axoneme components at the tips of flagella is consistent with a defect in axoneme/flagella length homeostasis (Qin *et al.*, 2004; Absalon *et al.*, 2008; Meng and Pan, 2016).

Microtubule staining also appears altered in Nek8445 knockdown cells. Specifically, we noticed that staining of the ventral disc was often reduced while the axonemes of Nek8445 depleted cells appeared more clearly delineated with less background staining. Our initial immunofluorescence localization was performed with the anti-acetylated α-tubulin antibody 6-11B-1 which is known to recognizes K40 acetylation and stain all known microtubule-based structures in *Giardia* (Nohynkova *et al.*, 2006; Halpern *et al.*, 2017). The altered staining indicates that either acetylation levels or microtubule array organization were perturbed by Nek8445 depletion. Microtubule acetylation is thought to soften microtubules so that they are more flexible and able to resist mechanical stress (reviewed in (Janke and Montagnac, 2017)). Notably, human Nek3 has been shown to regulate the levels of acetylated tubulin (Chang *et al.*, 2009). Therefore, we questioned whether Nek8445 has a role in regulating microtubule acetylation as this could potentially account for the observed defects in axoneme length and ventral disc staining.

To differentiate between changes in acetylation or overall microtubule organization, we co-stained cells with the anti-acetylated α-tubulin antibody 6-11B-1 and the anti-α-tubulin antibody DM1A. Total α-tubulin and acetylated tubulin localization appeared qualitatively similar in Nek8445 knockdown cells (Figure 4A). Immunoblotting with the two tubulin antibodies indicated that there is not a significant difference in the ratio of acetylated to non-acetylated α-tubulin when comparing control morpholino and Nek8445 knockdown cells (Figure 4D). This confirmed that that Nek8445 does not regulate acetylation. However, consistent with overall smaller cells that lack median bodies, we found that tubulin levels are approximately 25% lower in Nek8445 depleted cells compared to control morpholino treated cells.

### Nek8445 depleted cells have defective ventral disc organization

We previously reported that Nek8445 depleted cells were defective in their ability to attach to the sidewalls of culture tubes (Hennessey *et al.*, 2016). Attachment is largely attributed to the microtubule-based suction cup-like organelle known as the ventral adhesive disc (Dawson and Nosala, 2017). Since the ventral disc failed to stain with anti-tubulin antibodies in many of the Nek8445 depleted cells, we examined localization of the ventral disc marker δ-giardin to verify that a ventral disc was still present (Jenkins *et al.*, 2009; Halpern *et al.*, 2017). We did not find any cells lacking δ-giardin-3HA staining, indicating that the ventral disc was present (Figure 5). However, we observed abnormal disc organization that correlated with abnormal cell shape and flagella length. Of the Nek8445 depleted cells that consequently had short flagella, we observed 38% (n=108) of these cells to have an altered disc morphology compared to the control morpholino treated cells (n=63). Defects included a larger area devoid of δ-giardin staining in the center of the disc (bare area), frayed, and open disc conformations (Figure 5). Similar defects have been observed in cells depleted of disc associated proteins (DAPs) that have a structural role in the ventral disc (McInally *et al.*, 2019a); therefore, Nek8445 may have a role in regulating ventral disc assembly.

**Figure 5.**
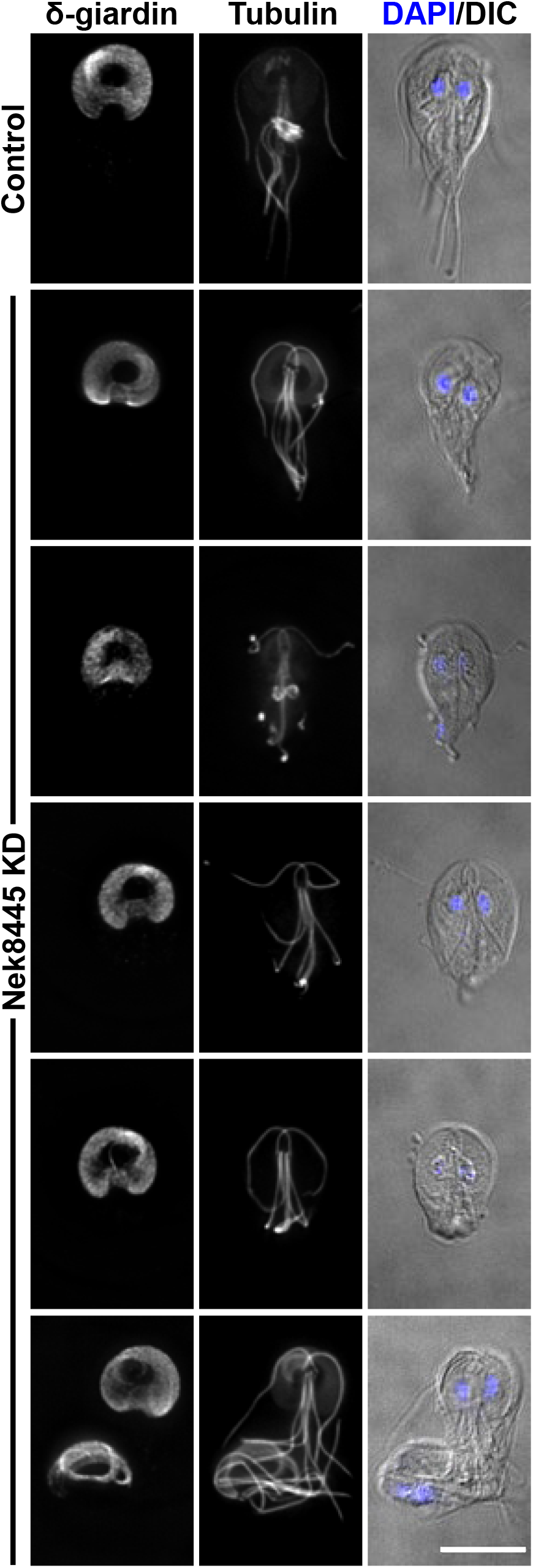
Nek8445 depleted cells have ventral disc organization defects. Control morpholino treated cells and Nek8445 knockdown cells stained for δ-giardin-3HA and tubulin (6-11B-1 antibody). Note the altered disc morphology such as increased width, larger bare area, and some striated staining of δ-giardin-3HA not observed in control cells. Of rounded Nek8445 depleted cells, 38% (n=108) had altered disc morphology not observed in the control morpholino treated cells (n=63). Scale bar = 5 μm.

### Nek8445 depleted cells have axoneme exit defects

In contrast to the typical half-pear shape of trophozoites, a subset of Nek8445 depleted cells were observed to have a rounded cell shape that lacked a portion of the posterior end of the cell. Tubulin staining (6-11B-1) indicated that the caudal flagella of these cells were either tucked under the cell body or the axonemes were trapped within the cell body, but this could not be resolved with light microscopy (Figure 4 and 5). To resolve whether the axonemes were trapped in the trophozoite cell body we turned to Scanning Electron Microscopy (SEM). Control morpholino (n=26) and translation blocking anti-Nek8445 treated cells (n=50) were processed for SEM 12 h after morpholino electroporation. As observed by light microscopy, a subset of the Nek8445 depleted cells were much smaller and had short flagella (Figure 6). While the anterior, ventral and posterolateral flagella were visible, the caudal flagella were frequently missing. In some cases, we could see abnormal membrane protrusions near where the caudal flagella were expected to exit, with successful exit from an atypical position (Figure 6 Arrowhead). The defect in axoneme exit site specification is expected to contribute to the cytokinesis defects observed with Nek8445 depletion as the flagella must be properly oriented in order to collaboratively generate forces for cytokinesis (Mariante *et al.*, 2005; Paredez *et al.*, 2011; Hardin *et al.*, 2017).

**Figure 6.**
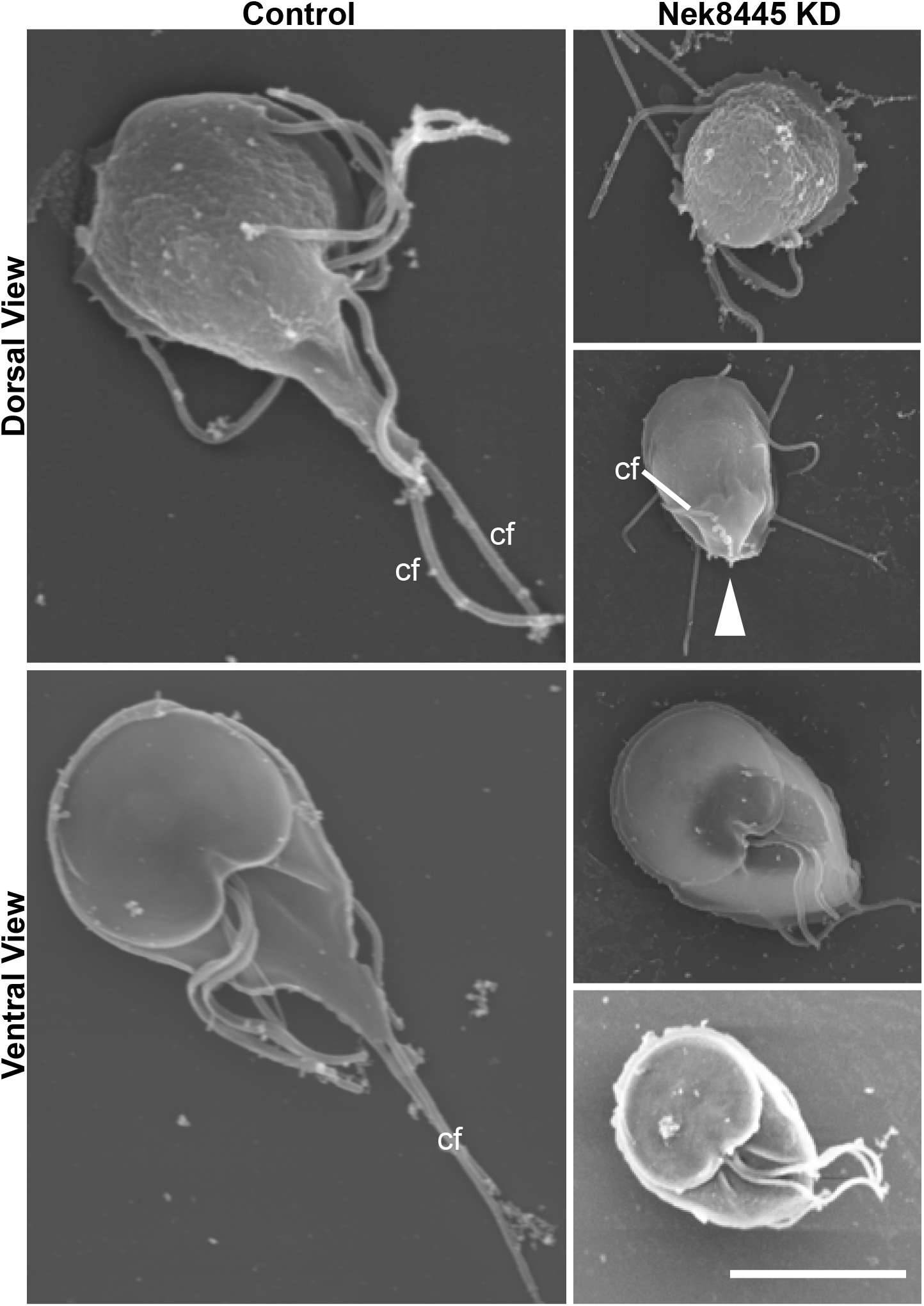
SEM analysis reveals Nek8445 depleted cells have caudal axoneme exit defects. Control morpholino and Nek8445 knockdown cells processed for scanning electron microscopy twelve hours after morpholino treatment. Note the shorter cell length and absence of caudal flagella (cf) at the anterior apex of the cell. White arrowhead indicates an abnormal membrane protrusion associated with a caudal flagellum that eventually exits the cell body at an abnormal position. Scale bar = 5 μm.

The defective axoneme exit defect provided us with the opportunity to investigate whether the exit site is pre-determined or established after the axonemes are in position. Unlike other eukaryotes where basal body appendages known as transition fibers dock the basal body to the plasma membrane (Goncalves and Pelletier, 2017), *Giardia* basal bodies are positioned many microns from where the cytoplasmic axoneme exits the cell body to become sheathed by plasma membrane. *Giardia* lacks all known transition zone proteins, so how the axoneme exit site is specified remains an open question (Avidor-Reiss & Leroux, 2015; Barker, Renzaglia, Fry, & Dawe, 2014). A recent study of *Giardia* intraflagellar transport (IFT) suggests the exit sites are pore-like diffusion barriers that regulate IFT (McInally *et al.*, 2019b). The Hehl lab, University of Zurich, identified GL50803_8855 as a protein that forms a ring at the axoneme exit site, where the axonemes transition from cytoplasmic to membrane sheathed (GiardiaDB). We endogenously tagged Exit Site Protein GL50803_8855 (ESP8855) with a triple HA tag to test whether the exit site was altered in the Nek8445 knockdown cells. As previously reported, we observed distinct localization of ESP8855-HA to the exit site of each axoneme set in cells treated with the control morpholino (n=23) (Figure 7A). In Nek8445 knockdown cells, ESP8855 localization varied. In cells where the caudal axoneme exited the cell body, ESP8855-HA was present where axonemes met the plasma membrane (Figure 7B). This included several cells with abnormally oriented caudal cytoplasmic axonemes (6 of 58) that mispositioned their caudal exit sites (Figure 7C). However, ESP8855-HA did not mark the posterior exit site in 59% (n=58) of rounded cells with short caudal axonemes that in some cases were too short to reach the cell cortex (Figure 7D). Together we interpret these findings to indicate that the exit site is specified upon axoneme contact with the plasma membrane.

**Figure 7.**
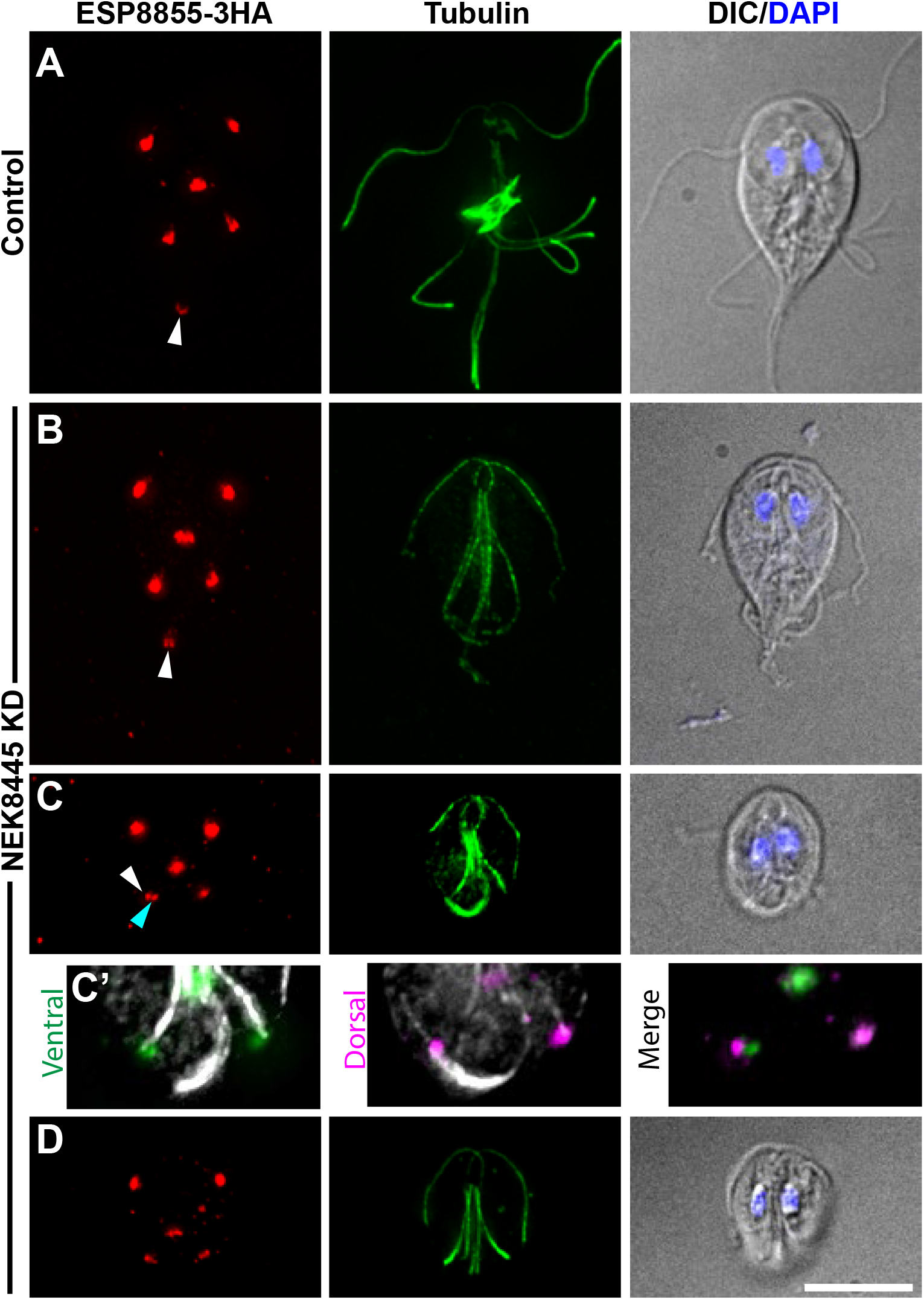
Nek8445 depleted cells misposition or lack exit sites. Control morpholino and Nek8445 knockdown cells were stained for tubulin (6-11B-1 antibody, green) and ESP8855-3HA (red). White arrowheads mark the caudal axoneme exit site and are oriented with the trajectory of the exiting axoneme. (A) Typical exit site organization in control cells. (B) Knockdown cell with typical teardrop shape retains exit sites at the expected position. (C) Mispositioned caudal axoneme exit site observed in 10% of rounded knockdown cells (n=58). Note these images are maximum projections, the caudal axoneme exit site in this cell is clearly positioned above the adjacent posterolateral axoneme exit site (cyan arrowhead) when viewed in the 3D image stack. (C’) Shows maximum projections of the ventral and dorsal halves of the Z-stack to show that these exit sites are displaced in Z similar to the cell with the arrowhead in Figure 6. (D) Complete lack of a caudal axoneme exit site is observed in 59% of rounded knockdown cells (n=58). Scale bar = 5 μm.

### Super-Resolution Imaging reveals Nek8445 depleted cells lack a funis

Nek8445 depletion alters the lengths of all flagella, but the caudal axonemes were specifically susceptible to being trapped in the cell body without an exit site. Many of the trapped axonemes are long enough to exit the cell body; therefore, the failure to exit cannot simply be attributed to length. The caudal cytoplasmic axonemes make the longest run through the cell body and are uniquely sandwiched between the two halves of the funis (see Figure 1). Since the cytoplasmic caudal axonemes often stained with higher contrast than in controls, and the exit sites were sometimes mispositioned (Figure 4, 5, 6, 7), we questioned whether the funis was lost or disrupted as a consequence of Nek8445 depletion. The funis is difficult to observe with standard resolution microscopy because it has overlapping localization with the caudal cytoplasmic axonemes and spacing of the rib-like microtubules is near the resolution limit of standard light microscopy. We turned to structured illumination super-resolution microscopy to compare control and Nek8445 depleted cells. We were able to resolve the funis in every control morpholino treated cell examined (n=12) while none of the rounded Nek8445 knockdown cells (n=12) had any vestige of a funis (Figure 8). The lack of a funis is consistent with the abnormal posterolateral axoneme/flagella positioning observed in Nek8445 knockdown cells. Previously observed connections between the funis microtubules and the posterolateral cytoplasmic axonemes were anticipated to be important for caudal cell movement (Benchimol *et al.*, 2004); however, the funis also appears to have a role in stabilizing the position of the posterolateral axonemes. Another unique feature of the rounded cells is that we observed several cells with frayed axoneme tips, further suggesting an axoneme stability defect in Nek8445 depleted cells (Wloga *et al.*, 2006; Bower *et al.*, 2013; Meng and Pan, 2016). The correlation of round cell shapes, mis-positioned axonemes/flagella, defective axoneme exit, and the lack of a funis is consistent with the funis having a role in establishing *Giardia* cell shape and guiding caudal axoneme exit. Together this study illustrates the overall importance of Nek8445 in regulating the microtubule cytoskeleton of *Giardia.*

**Figure 8.**
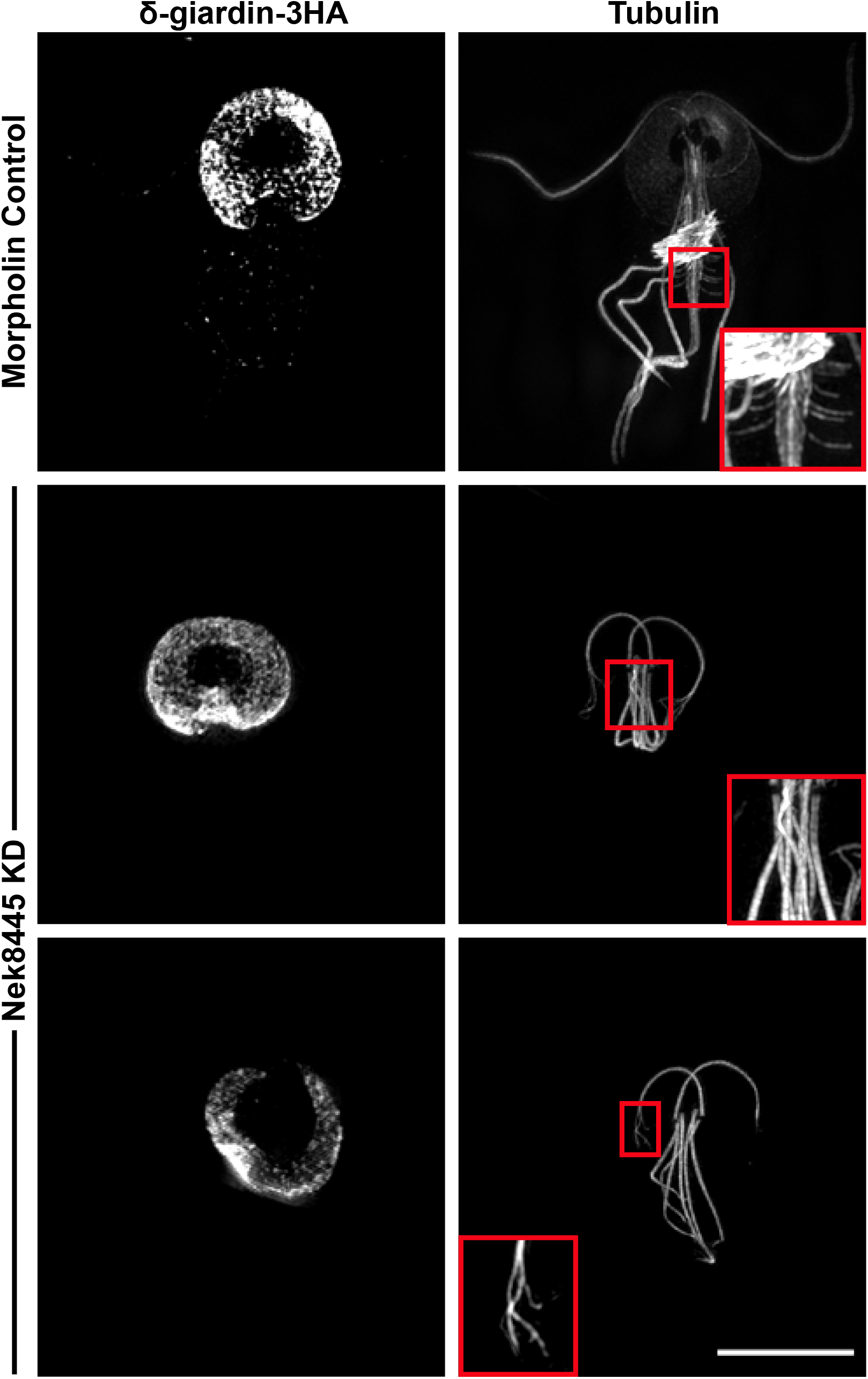
Structured Illumination super-resolution imaging of δ-giardin-3HA and tubulin (6-11B-1 antibody) in morpholino treated cells. Nek8445 depleted cells lack median bodies, a funis, and the axonemes have a fraying defect. Control inset shows magnified view of funis microtubules. Middle inset shows that the Nek8445 depleted cells lack both a median body and funis. The bottom image shows an example of a Nek8445 depleted cell with abnormal disc morphology and splayed axoneme tips. Scale bar = 5 μm.

## Discussion

We set out to study the primary cause of cytokinesis defects in Nek8445 depleted cells. Recent advances in live cell imaging have made it possible to visualize *Giardia* mitosis and cytokinesis with fantastic spatial-temporal resolution providing insights into *Giardia’s* myosin-independent mechanism of cytokinesis (Hardin *et al.*, 2017). Characterization of the Nek8445 knockdown phenotypes illustrates a requirement for Nek8445 in coordinating microtubule organization. This study reinforces the view that the microtubule cytoskeleton has expanded function for *Giardia* cell division, which includes force generation necessary for cytokinesis (Benchimol, 2004; Correa and Benchimol, 2006; Hardin *et al.*, 2017).

Our working model for myosin-independent cytokinesis (Hardin *et al.*, 2017), posits that at the onset of cytokinesis, the caudal cytoplasmic axonemes, each connecting to a daughter ventral disc, orients the daughter discs so that the anterior-most regions face away from one another. Major re-organization of the flagella occur as the cells initiate cytokinesis such that each daughter cell will inherit four full length flagella and grow four new flagella (Nohynkova *et al.*, 2006; Sagolla *et al.*, 2006; Tumova *et al.*, 2007; Hardin *et al.*, 2017). As the furrow expands, the newly duplicated axonemes, which are completely contained within the cytoplasm of the dividing cell, are used as tracks to guide additional membrane into the cleavage furrow (Hardin *et al.*, 2017). Ultimately, the four mature flagella assume the position and identities of the anterior and caudal flagella (Nohynkova *et al.*, 2006; Hardin *et al.*, 2017). Power strokes from the anterior flagella drive the two daughter cells in opposite directions generating the membrane tension required for division (Zhang and Robinson, 2005; Ralston *et al.*, 2006; Poirier *et al.*, 2012; Hardin *et al.*, 2017).

The observation that Nek8445 knockdown cells with typical cell shape and organization fail cytokinesis (Figure 2), suggests that the flagella may have improper function prior to the extensive shortening observed in rounded cells. Cytokinesis in these cells appears to fail because the caudal axonemes are not able to orient the daughter disks in opposition to each other (Figure 2 and Video S1). This re-orientation requires both coordinated force generation and the delivery of plasma membrane to provide slack for the discs to move away from each other (Hardin *et al.*, 2017). The similarity between the failed cytokinesis phenotype of Brefeldin A treatment and Nek8445 depleted cells is striking (Hardin *et al.*, 2017), which leads us to suspect that membrane trafficking could be impaired.

While most 12 h post knockdown cells had typical cell shape and organization as assessed with DIC optics, tubulin staining revealed that a significant portion of cells (~45%) lack median bodies. The lack of median bodies is an indicator of reduced tubulin reservoirs. During mitosis, microtubules flux from the median body, to the spindle, nascent disks, and finally to the nascent axonemes (Nohynkova *et al.*, 2006; Hardin *et al.*, 2017). Growth of new axonemes could be impaired due to a limited tubulin supply. Importantly, the nascent axonemes point into the cleavage furrow during cytokinesis and are used to direct new membrane into the furrow. Thus, short or improperly organized nascent axonemes could potentially reduce the availability of membrane for furrowing. Due to the lack of robust cell synchronization and the very short window during which cells use these tracks to deliver membrane, it is technically challenging to find knockdown cells actively dividing in order to examine the positioning and length of the nascent axonemes.

Short flagella may indicate a defect in IFT. Such a role for Nek8445 would not be surprising as Nek family kinases are known to regulate IFT (Lin *et al.*, 2003; Surpili *et al.*, 2003; Wloga *et al.*, 2006; Meng and Pan, 2016). We did not measure axoneme or flagella growth rates, but the presence of short flagella with swollen tips is consistent with an IFT defect. Indeed, the short flagella phenotype observed in Nek8445 depleted cells is reminiscent of the *Giardia* Kinesin-2 mutants which impair IFT (Hoeng *et al.*, 2008; Carpenter and Cande, 2009).

The caudal axoneme exit defect provided a unique opportunity to probe the mechanism of exit site specification/flagella pore formation in *Giardia.* ESP8855 marks axoneme exit sites. When Nek8445 was depleted, cells with extremely short cytoplasmic axonemes were observed to lack ESP8855 staining at the posterior apex of the cell, which is where the caudal axonemes exit in wild-type interphase cells. Additionally, some cells had mispositioned posterolateral and caudal axonemes with improperly positioned exit sites (Figure 6 and 7C). These findings are consistent with axoneme contact with the plasma membrane being the specifier for exit site/flagella pore formation rather than the axoneme being targeted to a pre-determined position on the membrane. Since only the caudal flagella were observed to lack exit sites, it is likely a secondary defect of Nek8445 depletion.

The caudal cytoplasmic axonemes make the longest run through the cell body and are unique in their association with the funis, a ribbon of microtubules that runs along the length the cytoplasmic axoneme of each caudal flagellum (Benchimol *et al.*, 2004). The ribbon originates near the basal bodies and as it progresses toward the posterior, individual microtubules splay out to form rib-like structures that match the shape of the cell (Figure 1, 8) (Benchimol *et al.*, 2004). A defect in axoneme guidance or stabilization could lead to defects in specifying the exit site if for example the axoneme must remain in place for the exit site to be assembled. Indeed, the axoneme exit defect was specifically associated with rounded Nek8445 depleted cells which we found also lack funis microtubules (Fig 8). This suggests that the funis is responsible for establishing posterior cell morphology and additionally acts as a scaffold for guiding caudal axoneme exit.

Although many Nek8445 depleted cells had short flagella we interestingly observed other cells with longer cytoplasmic axonemes trapped within the cell. This is consistent with the idea that *Giardia* can employ IFT independent cytoplasmic axoneme growth as cytoplasmic axonemes do not necessarily need active IFT for growth (Briggs *et al.*, 2004; McInally *et al.*, 2019b). The importance of this observation remains to be determined. IFT has been assessed in interphase *Giardia,* where it has been shown that IFT train assembly occurs at the flagella pore/axoneme exit site (McInally *et al.*, 2019b). Regulation of new axoneme growth in mitotic cells remains unstudied. IFT-independent growth is likely important for the growth of nascent axonemes, which begin completely within the cell cytoplasm. We previously noted that after cytokinesis the ventral flagella, which have the shortest run through the cell body, re-grow more quickly than the posterolateral pair (Hardin *et al.*, 2017). Whether this is due to regulated growth or the fact that the ventral axoneme would be the first to initiate an exit site and establish regulated IFT remains to be determined.

For an otherwise minimalistic parasite, the hugely expanded family of Nek kinases is intriguing. *Giardia* has 198 members compared with just one non-essential Nek Kinase in yeast, seven in *Arabidopsis,* 11 in humans, 13 in *Chlamydomonas,* and 39 in *Tetrahymena* (Wloga *et al.*, 2006; Manning *et al.*, 2011; Takatani *et al.*, 2015). Each of the three *Giardia* Nek kinases that have been studied appear to regulate different cellular processes. Depletion studies of *Gl*Nek1 and *Gl*Nek2 have not been performed so whether they are required for cell division remains to be determined. However, localization studies did reveal that these proteins re-localized during mitosis. In general, Nek kinases are activated during the G2/M transition by mitotic kinases (Fry *et al.*, 2017), but Nek kinases have also been shown to function during interphase (Chang *et al.*, 2009; Cohen *et al.*, 2013; Govindaraghavan *et al.*, 2014). While we cannot rule out interphase function, re-localization of Nek8445, *Gl*Nek1, and *Gl*Nek2 during mitosis suggests these kinases likely regulate the microtubule cytoskeleton in a cell cycle dependent manner (Smith *et al.*, 2012). The Anaphase Promoting Complex (APC), which canonically regulates cell cycle events through proteolytic degradation, has not been identified in *Giardia* (Gourguechon *et al.*, 2013). It has been speculated that kinases could play an expanded role in regulating cell cycle related process (Gourguechon *et al.*, 2013); the expanded Nek kinase family potentially fills this role. Future studies aimed at identifying pathways regulated by additional Nek kinases will be needed to further support this hypothesis.

Given the total number of Nek kinases in *Giardia,* we anticipated that each kinase would have a limited role in cytoskeletal regulation, yet Nek8445 depletion disrupts nearly all of the major microtubule structures in *Giardia.* The exception is that Nek8445 does not appear to regulate centrosome separation or spindle function as mitosis appears to progress normally. The other microtubule defects could result from a role for Nek8445 in regulating a key microtubule binding protein with a role in regulating *Giardia’s* microtubule-based structures. For instance, EB1 has important roles in regulating microtubule dynamics and flagella length (Pedersen *et al.*, 2005; Schroder *et al.*, 2007). Several studies have localized Nek Kinases to centrosomes, the tips of microtubules and axonemes and implicated a relationship with EB1 (Mahjoub *et al.*, 2004; O’Regan and Fry, 2009; Govindaraghavan *et al.*, 2014; Prosser *et al.*, 2015). The relationship between Nek kinases and EB1 is not fully characterized, but *Aspergillus nimA* has been shown to require EB1 for its localization (Govindaraghavan *et al.*, 2014). Intriguingly, EB1 knockdown in *Giardia* has been shown to reduce median body volume, flagella length, and cause loss of axoneme central pairs (Kim and Park, 2019). Although less severe, these phenotypes are similar to what we observe in Nek8445 depleted cells. It will be interesting to identify Nek8445 phosphorylation targets to gain insight into microtubule array regulation in *Giardia.*

Whether many other Nek kinases play as critical a role as Nek8445 in cell division and microtubule array biogenesis remains to be determined, but the presence of a small gatekeeper residue in Nek8445 makes this protein kinase a particularly attractive drug target as this structural feature renders this kinase sensitive to bumped kinase inhibitors. Importantly, we have already shown this class of inhibitor is effective at killing parasites resistant to metronidazole (Hennessey *et al.*, 2016), the frontline treatment for giardiasis.

## Acknowledgements

We thank Christine Wong for technical assistance and Wai Pang for help with the SEM. We thank Elizabeth Thomas and Melissa Steele-Ogus for help with editing and preparing figures. This work was supported by NIH award R01AI110708 to A.R.P and R21AI127493 to E.A.M. The funders had no role in study design, data collection and interpretation, or the decision to submit the work for publication.

---

**Figure S1** Phenotype of Nek8445 knockdown cells at 48 hours post morpholino treatment. Nek8445-HA stained in red, tubulin in green (6-11B-1 antibody), and nuclei in blue. Asterisks mark six cells with abnormal nuclear number and altered cell shape. These defects correlate with reduced Nek8445 levels as indicated by staining intensity. The presence of cells with more than 4 nuclei indicates that although these cells failed cytokinesis they continue to proceed through mitosis. Scale bar= 10 μm.

**Figure S2** Start point for long-term DIC movies. (A) Few cells are able to attach to the cover glass at 22 hours post Nek8445 knockdown. (B) Quantification of multinucleate (>2 nuclei) cells that are either in the process of dividing or have failed to divide (Control Morpholino=6.4%, KD=51.9%; n=2400 for each treatment). (C) Control morpholino treated cells at 12 hours post electroporation. (D) Nek8445 antisense morpholino-treated cells 12 hours post knockdown. Scale bar= 5 μm.

**Figure S3.** Rab11 and Nek8445 localization throughout the cell cycle. HA-Rab11 in cyan, Nek8445-mNG in red, and tubulin (6-11B-1 antibody) in green. During mitosis Nek8445 is associated with the nuclear envelope and loads onto cytoplasmic axonemes. HA-Rab11 also loads onto cytoplasmic axonemes at specific times during mitosis, but Rab11 and Nek8445 are not frequently observed to co-localize. White arrowhead shows an example of HA-Rab11 and Nek8445-mNG co-localization on the cytoplasmic axonemes of the posterolateral flagella. Scale bar= 5 μm.

**Video S1** Control morpholino and translation blocking Nek8445 morpholino treated cells undergoing mitosis and cytokinesis. Note that the videos have been synchronized by the start of mitosis as indicated by anterior flagella re-organization.

**Video S2** Live imaging of Nek8445-mNG during mitosis and cytokinesis.

## References

Absalon, S., Blisnick, T., Bonhivers, M., Kohl, L., Cayet, N., Toutirais, G., Buisson, J., Robinson, D., and Bastin, P. (2008). Flagellum Elongation Is Required for Correct Structure, Orientation and Function of the Flagellar Pocket in Trypanosoma Brucei. J Cell Sci 121, 3704–3716.

Ansell, B.R., McConville, M.J., Ma’ayeh, S.Y., Dagley, M.J., Gasser, R.B., Svard, S.G., and Jex, A.R. (2015). Drug Resistance in Giardia Duodenalis. Biotechnology advances 33, 888–901.

Benchimol, M. (2004). Participation of the Adhesive Disc During Karyokinesis in Giardia Lamblia. Biology of the cell/ under the auspices of the European Cell Biology Organization 96, 291–301.

Benchimol, M., Piva, B., Campanati, L., and de Souza, W. (2004). Visualization of the Funis of Giardia Lamblia by High-Resolution Field Emission Scanning Electron Microscopy--New Insights. J Struct Biol 147, 102–115.

Bower, R., Tritschler, D., Vanderwaal, K., Perrone, C.A., Mueller, J., Fox, L., Sale, W.S., and Porter, M.E. (2013). The N-Drc Forms a Conserved Biochemical Complex That Maintains Outer Doublet Alignment and Limits Microtubule Sliding in Motile Axonemes. Mol Biol Cell 24, 1134–1152.

Bradley, B.A., and Quarmby, L.M. (2005). A Nima-Related Kinase, Cnk2p, Regulates Both Flagellar Length and Cell Size in Chlamydomonas. J Cell Sci 118, 3317–3326.

Briggs, L.J., Davidge, J.A., Wickstead, B., Ginger, M.L., and Gull, K. (2004). More Than One Way to Build a Flagellum: Comparative Genomics of Parasitic Protozoa. Current biology: CB 14, R611–612.

Brown, J.R., Schwartz, C.L., Heumann, J.M., Dawson, S.C., and Hoenger, A. (2016). A Detailed Look at the Cytoskeletal Architecture of the Giardia Lamblia Ventral Disc. J Struct Biol 194, 38–48.

Carpenter, M.L., and Cande, W.Z. (2009). Using Morpholinos for Gene Knockdown in Giardia Intestinalis. Eukaryot Cell 8, 916–919.

Carvalho, K.P., and Monteiro-Leal, L.H. (2004). The Caudal Complex of Giardia Lamblia and Its Relation to Motility. Experimental parasitology 108,154–162.

Chang, J., Baloh, R.H., and Milbrandt, J. (2009). The Nima-Family Kinase Nek3 Regulates Microtubule Acetylation in Neurons. J Cell Sci 122, 2274–2282.

Chen, C.T., and Gubbels, M.J. (2013). The Toxoplasma Gondii Centrosome Is the Platform for Internal Daughter Budding as Revealed by a Nek1 Kinase Mutant. Journal of cell science 126, 3344–3355.

Cohen, S., Aizer, A., Shav-Tal, Y., Yanai, A., and Motro, B. (2013). Nek7 Kinase Accelerates Microtubule Dynamic Instability. Biochimica et biophysica acta 1833,1104–1113.

Correa, G., and Benchimol, M. (2006). Giardia Lamblia Behavior under Cytochalasins Treatment. Parasitology research 98, 250–256.

Dawson, S.C., and Nosala, C. (2017). The Ventral Disc Is a Flexible Microtubule Organelle That Depends on Domed Ultrastructure for Functional Attachment of Giardia Lamblia. bioRxiv.

Faragher, A.J., and Fry, A.M. (2003). Nek2a Kinase Stimulates Centrosome Disjunction and Is Required for Formation of Bipolar Mitotic Spindles. Mol Biol Cell 14, 2876–2889.

Fry, A.M., Bayliss, R., and Roig, J. (2017). Mitotic Regulation by Nek Kinase Networks. Frontiers in cell and developmental biology 5,102.

Goncalves, J., and Pelletier, L. (2017). The Ciliary Transition Zone: Finding the Pieces and Assembling the Gate. Molecules and cells 40, 243–253.

Gourguechon, S., and Cande, W.Z. (2011). Rapid Tagging and Integration of Genes in Giardia Intestinalis. Eukaryot Cell 10,142–145.

Gourguechon, S., Holt, L.J., and Cande, W.Z. (2013). The Giardia Cell Cycle Progresses Independently of the Anaphase-Promoting Complex. Journal of Cell Science 126, 2246–2255.

Govindaraghavan, M., McGuire Anglin, S.L., Shen, K.F., Shukla, N., De Souza, C.P., and Osmani, S.A. (2014). Identification of Interphase Functions for the Nima Kinase Involving Microtubules and the Escrt Pathway. PLoS genetics 10, e1004248.

Halpern, A.R., Alas, G.C.M., Chozinski, T.J., Paredez, A.R., and Vaughan, J.C. (2017). Hybrid Structured Illumination Expansion Microscopy Reveals Microbial Cytoskeleton Organization. ACS nano 11, 12677–12686.

Hansen, W.R., Tulyathan, O., Dawson, S.C., Cande, W.Z., and Fletcher, D.A. (2006). Giardia Lamblia Attachment Force Is Insensitive to Surface Treatments. Eukaryot. Cell 5, 781–783.

Hardin, W.R., Li, R., Xu, J., Shelton, A.M., Alas, G.C.M., Minin, V.N., and Paredez, A.R. (2017). Myosin-Independent Cytokinesis in Giardia Utilizes Flagella to Coordinate Force Generation and Direct Membrane Trafficking. Proc Natl Acad Sci USA.

Hennessey, K.M., Smith, T.R., Xu, J.W., Alas, G.C., Ojo, K.K., Merritt, E.A., and Paredez, A.R. (2016). Identification and Validation of Small-Gatekeeper Kinases as Drug Targets in Giardia Lamblia. PLoS neglected tropical diseases 10, e0005107.

Hoeng, J.C., Dawson, S.C., House, S.A., Sagolla, M.S., Pham, J.K., Mancuso, J.J., Lowe, J., and Cande, W.Z. (2008). High-Resolution Crystal Structure and in Vivo Function of a Kinesin-2

Erlandsen, S.L., Russo, A.P., and Turner, J.N. (2004). Evidence for Adhesive Activity of the Ventrolateral Flange in Giardia Lamblia. The Journal of eukaryotic microbiology 51, 73–80.

Homologue in Giardia Intestinalis. Mol Biol Cell 19, 3124–3137.

Holberton, D.V. (1973). Fine Structure of the Ventral Disk Apparatus and the Mechanism of Attachment in the Flagellate Giardia Muris. J Cell Sci 13,11–41.

Horlock-Roberts, K., Reaume, C., Dayer, G., Ouellet, C., Cook, N., and Yee, J. (2017). Drug-Free Approach to Study the Unusual Cell Cycle of Giardia Intestinalis. mSphere 2.

House, S.A., Richter, D.J., Pham, J.K., and Dawson, S.C. (2011). Giardia Flagellar Motility Is Not Directly Required to Maintain Attachment to Surfaces. PLoS Pathog 7, e1002167.

Janke, C., and Montagnac, G. (2017). Causes and Consequences of Microtubule Acetylation. Current biology: CB 27, R1287–r1292.

Jenkins, M.C., O’Brien, C.N., Murphy, C., Schwarz, R., Miska, K., Rosenthal, B., and Trout, J.M. (2009). Antibodies to the Ventral Disc Protein Delta-Giardin Prevent in Vitro Binding of Giardia Lamblia Trophozoites. The Journal of parasitology 95, 895–899.

Keister, D.B. (1983). Axenic Culture of Giardia-Lamblia in Tyi-S-33 Medium Supplemented with Bile. Transactions of the Royal Society of Tropical Medicine and Hygiene 77, 487–488.

Keyloun, K.R., Reid, M.C., Choi, R., Song, Y., Fox, A.M.W., Hillesland, H.K., Zhang, Z., Vidadala, R., Merritt, E.A., Lau, A.O.T., Maly, D.J., Fan, E., Barrett, L.K., Van Voorhis, W.C., and Ojo, K.K. (2014). The Gatekeeper Residue and Beyond: Homologous Calcium-Dependent Protein Kinases as Drug Development Targets for Veterinarian Apicomplexa Parasites. Parasitology 141, 1499–1509.

Kim, J., and Park, S.J. (2019). Roles of End-Binding 1 Protein and Gamma-Tubulin Small Complex in Cytokinesis and Flagella Formation of Giardia Lamblia. Microbiology Open 8, e00748.

Krtkova, J., and Paredez, A.R. (2017). Use of Translation Blocking Morpholinos for Gene Knockdown in Giardia Lamblia. Methods in molecular biology (Clifton, N.J.) 1565, 123–140.

Lalle, M. (2010). Giardiasis in the Post Genomic Era: Treatment, Drug Resistance and Novel Therapeutic Perspectives. Infectious disorders drug targets 10, 283–294.

Lane, S., and Lloyd, D. (2002). Current Trends in Research into the Waterborne Parasite Giardia. Crit Rev Microbiol 28,123–147.

Lenaghan, S.C., Davis, C.A., Henson, W.R., Zhang, Z., and Zhang, M. (2011). High-Speed Microscopic Imaging of Flagella Motility and Swimming in Giardia Lamblia Trophozoites. Proc Natl Acad Sci U S A 108, E550–558.

Lin, F., Hiesberger, T., Cordes, K., Sinclair, A.M., Goldstein, L.S., Somlo, S., and Igarashi, P. (2003). Kidney-Specific Inactivation of the Kif3a Subunit of Kinesin-li Inhibits Renal Ciliogenesis and Produces Polycystic Kidney Disease. Proc Natl Acad Sci U S A 100, 5286–5291.

Mahjoub, M.R., Qasim Rasi, M., and Quarmby, L.M. (2004). A Nima-Related Kinase, Fa2p, Localizes to a Novel Site in the Proximal Cilia of Chlamydomonas and Mouse Kidney Cells. Mol Biol Cell 15, 5172–5186.

Manning, G., Reiner, D.S., Lauwaet, T., Dacre, M., Smith, A., Zhai, Y., Svard, S., and Gillin, F.D. (2011). The Minimal Kinome of Giardia Lamblia Illuminates Early Kinase Evolution and Unique Parasite Biology. Genome biology 12, R66.

Mariante, R.M., Vancini, R.G., Melo, A.L., and Benchimol, M. (2005). Giardia Lamblia: Evaluation of the in Vitro Effects of Nocodazole and Colchicine on Trophozoites. Experimental parasitology 110, 62–72.

McInally, S.G., and Dawson, S.C. (2016). Eight Unique Basal Bodies in the Multi-Flagellated Diplomonad Giardia Lamblia. Cilia 5, 21.

McInally, S.G., Hagen, K.D., Nosala, C., Williams, J., Nguyen, K., Booker, J., Jones, K., and Dawson, S.C. (2019a). Robust and Stable Transcriptional Repression in Giardia Using Crispri. Mol Biol Cell 30, 119–130.

McInally, S.G., Kondev, J., and Dawson, S.C. (2019b). Length-Dependent Disassembly Maintains Four Different Flagellar Lengths in Giardia. eLife 8.

Meng, D., and Pan, J. (2016). A Nima-Related Kinase, Cnk4, Regulates Ciliary Stability and Length. Mol Biol Cell 27, 838–847.

Nohynkova, E., Tumova, P., and Kulda, J. (2006). Cell Division of Giardia Intestinalis: Flagellar Developmental Cycle Involves Transformation and Exchange of Flagella between Mastigonts of a Diplomonad Cell. Eukaryot Cell 5, 753–761.

Nosala, C., Hagen, K.D., and Dawson, S.C. (2018). ‘Disc-O-Fever’: Getting Down with Giardia’s Groovy Microtubule Organelle. Trends Cell Biol 28, 99–112.

O’Connell, M.J., Krien, M.J., and Hunter, T. (2003). Never Say Never. The Nima-Related Protein Kinases in Mitotic Control. Trends Cell Biol 13, 221–228.

O’Regan, L., and Fry, A.M. (2009). The Nek6 and Nek7 Protein Kinases Are Required for Robust Mitotic Spindle Formation and Cytokinesis. Mol Cell Biol 29, 3975–3990.

Paredez, A.R., Assaf, Z.J., Sept, D., Timofejeva, L., Dawson, S.C., Wang, C.J., and Cande, W.Z. (2011). An Actin Cytoskeleton with Evolutionarily Conserved Functions in the Absence of Canonical Actin-Binding Proteins. Proc Natl Acad Sci U S A 108, 6151–6156.

Parker, J.D., Bradley, B.A., Mooers, A.O., and Quarmby, L.M. (2007). Phylogenetic Analysis of the Neks Reveals Early Diversification of Ciliary-Cell Cycle Kinases. PloS one 2, e1076.

Pedersen, L.B., Miller, M.S., Geimer, S., Leitch, J.M., Rosenbaum, J.L., and Cole, D.G. (2005). Chlamydomonas Ift172 Is Encoded by Fla11, Interacts with Creb1, and Regulates Ift at the Flagellar Tip. Current biology: CB 15, 262–266.

Poirier, C.C., Ng, W.P., Robinson, D.N., and Iglesias, P.A. (2012). Deconvolution of the Cellular Force-Generating Subsystems That Govern Cytokinesis Furrow Ingression. Pios Computational Biology 8.

Prosser, S.L., Sahota, N.K., Pelletier, L., Morrison, C.G., and Fry, A.M. (2015). Nek5 Promotes Centrosome Integrity in Interphase and Loss of Centrosome Cohesion in Mitosis. The Journal of cell biology 209, 339–348.

Qin, H., Diener, D.R., Geimer, S., Cole, D.G., and Rosenbaum, J.L. (2004). Intraflagellar Transport (Ift) Cargo: Ift Transports Flagellar Precursors to the Tip and Turnover Products to the Cell Body. The Journal of cell biology 164, 255–266.

Ralston, K.S., Lerner, A.G., Diener, D.R., and Hill, K.L. (2006). Flagellar Motility Contributes to Cytokinesis in Trypanosoma Brucei and Is Modulated by an Evolutionarily Conserved Dynein Regulatory System. Eukaryot. Cell 5, 696–711.

Rotella, D.P. (2012). Recent Results in Protein Kinase Inhibition for Tropical Diseases. Bioorganic & medicinal chemistry letters 22, 6788–6793.

Sagolla, M.S., Dawson, S.C., Mancuso, J.J., and Cande, W.Z. (2006). Three-Dimensional Analysis of Mitosis and Cytokinesis in the Binucleate Parasite Giardia Intestinalis. J Cell Sci 119, 4889–4900.

Schneider, C.A., Rasband, W.S., and Eliceiri, K.W. (2012). Nih Image to Imagej: 25 Years of Image Analysis. Nature methods 9, 671–675.

Schroder, J.M., Schneider, L., Christensen, S.T., and Pedersen, L.B. (2007). Eb1 Is Required for Primary Cilia Assembly in Fibroblasts. Current biology: CB 17,1134–1139.

Shaner, N.C., Lambert, G.G., Chammas, A., Ni, Y., Cranfill, P.J., Baird, M.A., Sell, B.R., Allen, J.R., Day, R.N., Israelsson, M., Davidson, M.W., and Wang, J. (2013). A Bright Monomeric Green Fluorescent Protein Derived from Branchiostoma Lanceolatum. Nature methods 10, 407–409.

Smith, A.J., Lauwaet, T., Davids, B.J., and Gillin, F.D. (2012). Giardia Lamblia Nek1 and Nek2 Kinases Affect Mitosis and Excystation. Int J Parasitol 42, 411–419.

Surpili, M.J., Delben, T.M., and Kobarg, J. (2003). Identification of Proteins That Interact with the Central Coiled-Coil Region of the Human Protein Kinase Nek1. Biochemistry-Us 42, 15369–15376.

Takatani, S., Otani, K., Kanazawa, M., Takahashi, T., and Motose, H. (2015). Structure, Function, and Evolution of Plant Nima-Related Kinases: Implication for Phosphorylation-Dependent Microtubule Regulation. Journal of plant research 128, 875–891.

Tumova, P., Kulda, J., and Nohynkova, E. (2007). Cell Division of Giardia Intestinalis: Assembly and Disassembly of the Adhesive Disc, and the Cytokinesis. Cell motility and the cytoskeleton 64, 288–298.

Wloga, D., Camba, A., Rogowski, K., Manning, G., Jerka-Dziadosz, M., and Gaertig, J. (2006). Members of the Nima-Related Kinase Family Promote Disassembly of Cilia by Multiple Mechanisms. Mol Biol Cell 17, 2799–2810.

Zhang, W., and Robinson, D.N. (2005). Balance of Actively Generated Contractile and Resistive Forces Controls Cytokinesis Dynamics. Proc Natl Acad Sci U S A 102, 7186–7191.

