## Supplementary figures and images for "Nek8445, a protein kinase required for microtubule regulation and cytokinesis in *Giardia lamblia*"

### Figure S1

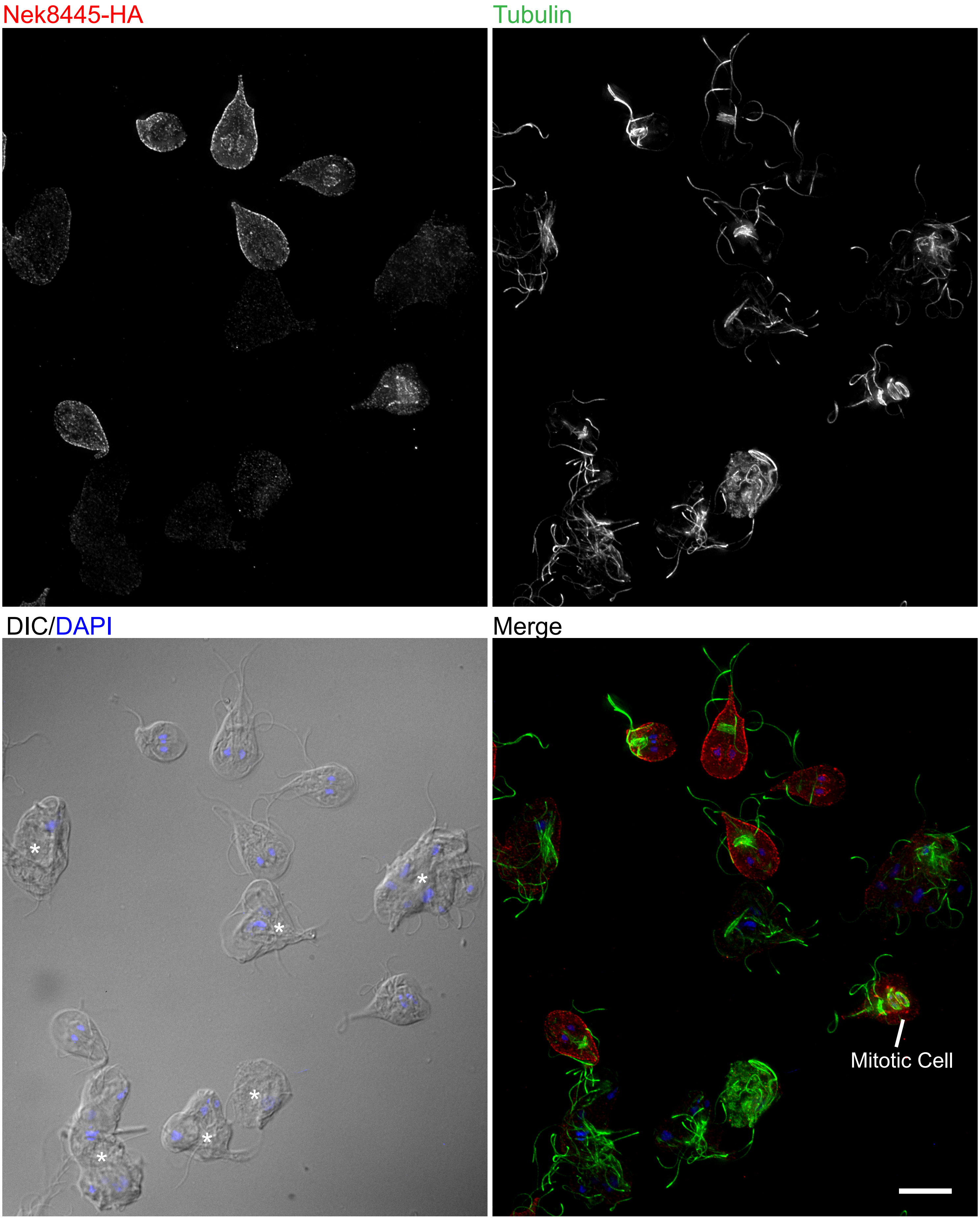

### Figure S2

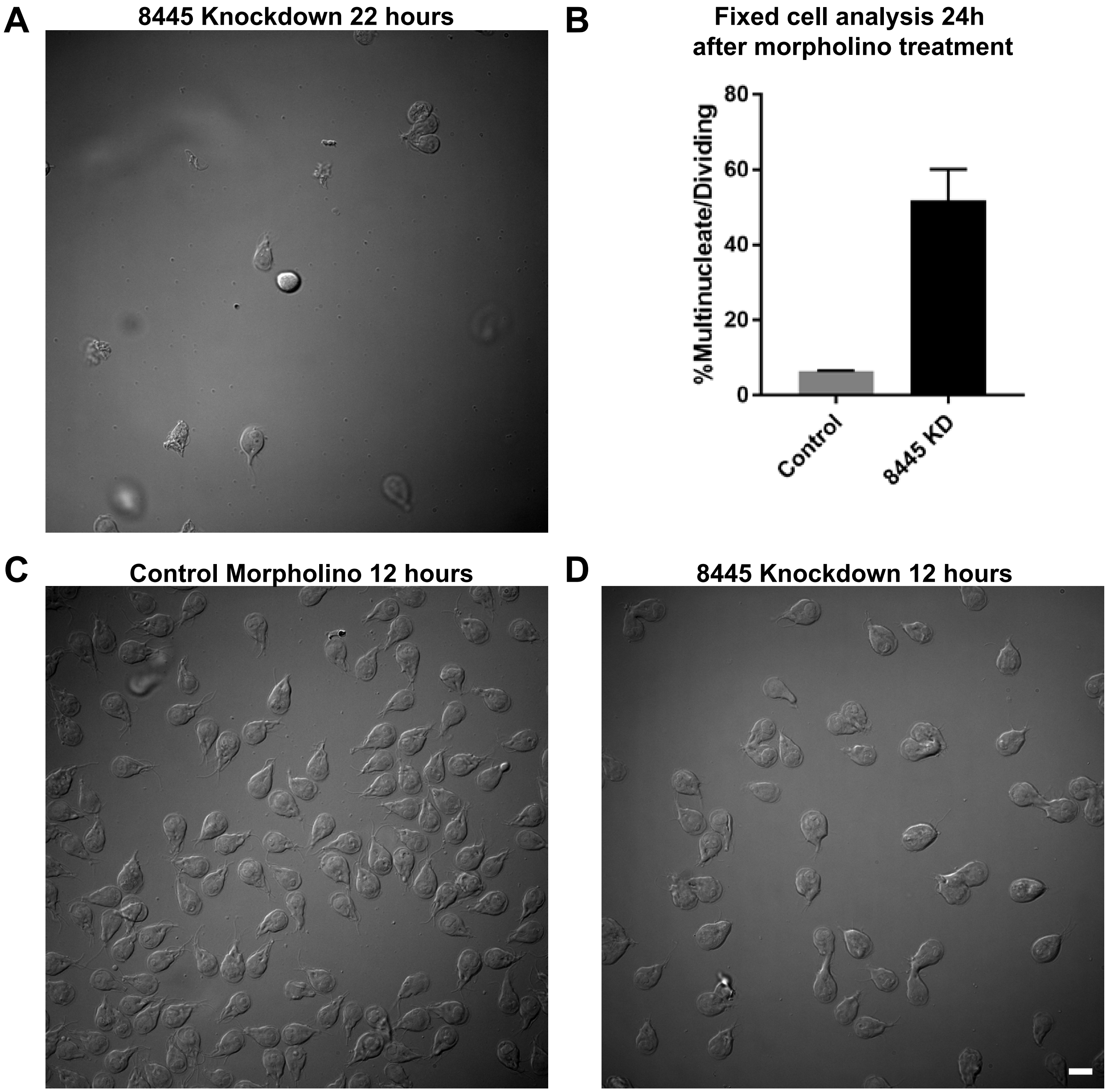

### Figure S3

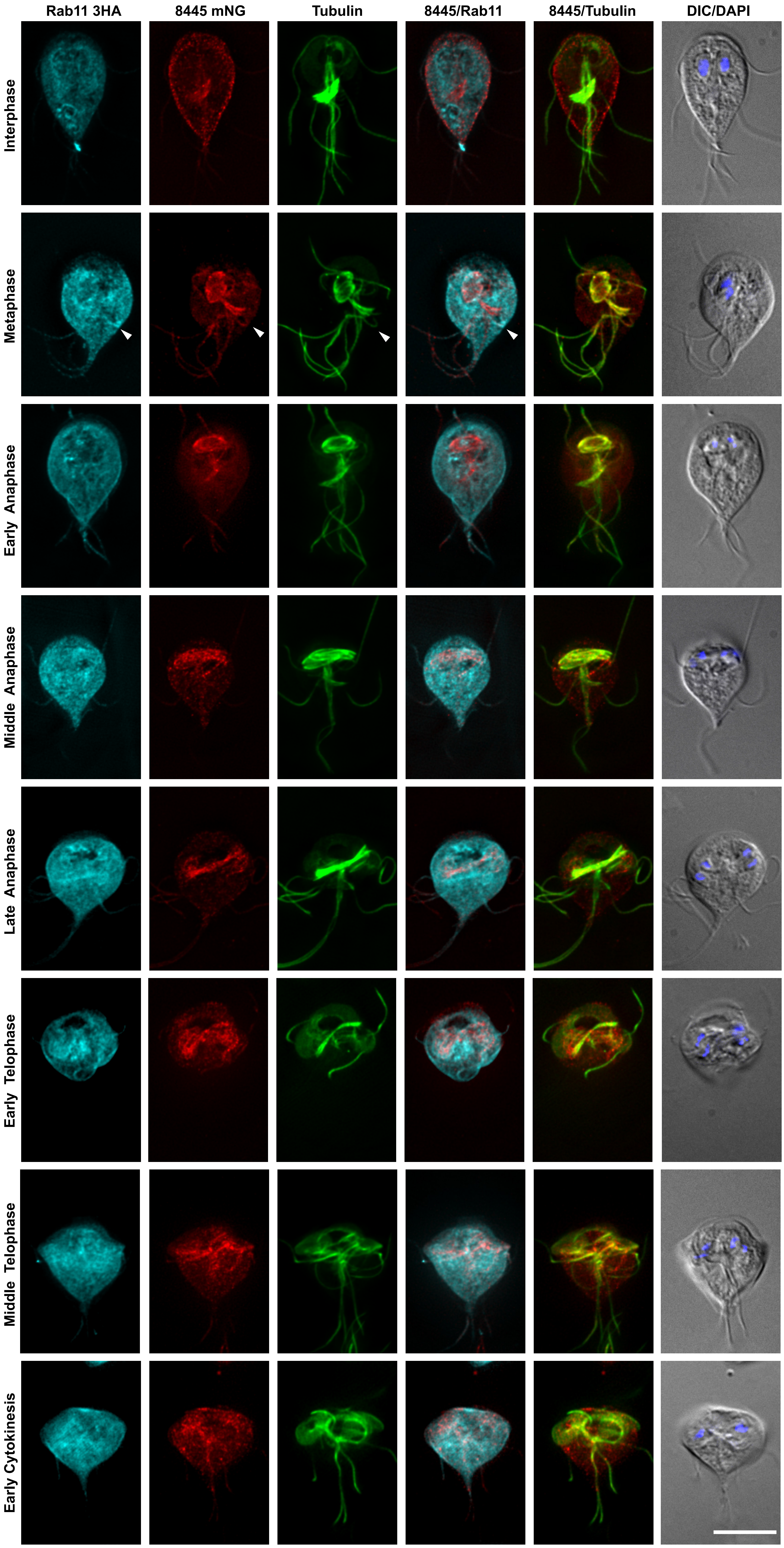
