## Supplementary material for "Nek8445, a protein kinase required for microtubule regulation and cytokinesis in *Giardia lamblia*": Table S1

| **PRIMER NAME** | **RESTRICTION ENZYME** | **5' TO 3' SEQUENCE** |
|---|---|---|
| 8445_VSVG_Neo R | AgeI | TTCAATATCAGTATAaccggtcttgttgttagtctgtagccat |
| 8445_VSVG_Neo F | NotI | tggagctccaccgcggtggcggccgcCATAATCTTGGCGTCCTTCG |
| 8445_mNG_Neo F | NotI | ctccaccgcggtggcggccgcCATAATCTTGGCGTCCTTCG |
| 8445_mNG_Neo R | BamHI | AACCGCCTCCGGATCCCTTGTTGTTAGTCTGTAGCC |
| 8855_3HA_Neo F | SpeI | cggtggcggccgctctagaactagtTGTTGAGGTATCTGGGGATATT |
| 8855_3HA_Neo R | AflII | gtcataaggatattccttaagAATATCAAAATCGTCGAAATCACC |
| pks_mNG_NEO_F | BamHI | cggccgctctagaactagtggatccGGAGGCGGTTCAGGCGGAGG TGGCTCTatggtgagcaagggcgagga |
| pks_mNG_NEO_R | EcoRI | Gcaacgatgatgcaaagaattcttacttgtacagctcgtccatg |
| Morpholino | | |
| 8445 | | CTGCCAGCTTCTCCGCAATACCCAT |

Table S1 Oligo sequences used in this study
